# Latent Epigenetic Programs in Müller Glia Contribute to Stress, Injury, and Disease Response in the Retina

**DOI:** 10.1101/2023.10.15.562396

**Authors:** Jackie L. Norrie, Marybeth Lupo, Abbas Shirinifard, Nadhir Djekidel, Cody Ramirez, Beisi Xu, Jacob M. Dundee, Michael A. Dyer

**Author notes:** Correspondence should be addressed to: Michael A. Dyer, PhD, Department of Developmental Neurobiology, St. Jude Children’s Research Hospital, 262 Danny Thomas Place, Memphis, TN 38105-3678, USA.

## Abstract

Previous studies have demonstrated the dynamic changes in chromatin structure during retinal development that correlate with changes in gene expression. However, a major limitation of those prior studies was the lack of cellular resolution. Here, we integrate single-cell (sc) RNA-seq and scATAC-seq with bulk retinal data sets to identify cell type–specific changes in the chromatin structure during development. Although most genes’ promoter activity is strongly correlated with chromatin accessibility, we discovered several hundred genes that were transcriptionally silent but had accessible chromatin at their promoters. Most of those silent/accessible gene promoters were in the Müller glial cells. The Müller cells are radial glia of the retina and perform a variety of essential functions to maintain retinal homeostasis and respond to stress, injury, or disease. The silent/accessible genes in Müller glia are enriched in pathways related to inflammation, angiogenesis, and other types of cell-cell signaling and were rapidly activated when we tested 15 different physiologically relevant conditions to mimic retinal stress, injury, or disease in human and murine retinae. We refer to these as “pliancy genes” because they allow the Müller glia to rapidly change their gene expression and cellular state in response to different types of retinal insults. The Müller glial cell pliancy program is established during development, and we demonstrate that pliancy genes are necessary and sufficient for regulating inflammation in the murine retina in vivo. In zebrafish, Müller glia can de-differentiate and form retinal progenitor cells that replace lost neurons. The pro-inflammatory pliancy gene cascade is not activated in zebrafish Müller glia following injury, and we propose a model in which species-specific pliancy programs underly the differential response to retinal damage in species that can regenerate retinal neurons (zebrafish) versus those that cannot (humans and mice).

## INTRODUCTION

Multipotent retinal progenitor cells produce 6 classes of neurons (rods, cones, bipolar cells, amacrine cells, horizontal cells, and ganglion cells) and 1 type of radial glia (Müller glia) in an evolutionarily conserved birth order during development [1, 2]. The Müller glia are one of the last cell types born during retinogenesis; their nuclei are centrally located, with cell bodies that span the entire thickness of the retina [3]. Müller glial cell processes surround the retinal neurons, and their apical microvilli form the outer limiting membrane on the apical surface of the retina [4, 5]. On the basal surface, Müller glial cell end feet contact the vitreous body and maintain K+ homeostasis in the retina. Beyond this spatial buffering of ions, Müller glia contribute to retinal homeostasis by recycling neurotransmitters, preventing glutamate toxicity, participating in the visual cycle, and serving as a source of nutrients [6–9]. In addition to retinal neurons, the Müller glia ensheath the retinal vasculature to form the blood-retinal barrier and are an important source of angiogenic factors [10, 11]. Taken together, decades of research have shown that Müller glia play multiple roles in maintaining retinal homeostasis and visual function.

The Müller glial cell response to stress, injury, and disease has diverged during evolution [12]. In fish and some amphibians, they de-differentiate into retinal progenitor cells, which then go on to replace the neurons that were lost during injury [12, 13]. In the mammalian retina, stress response pathways are activated in Müller glia to protect the retinal neurons, but they do not dedifferentiate into retinal progenitor cells or regenerate retinal neurons [14, 15]. The divergent molecular pathways that contribute to Müller glial cell regeneration versus stress response are the subject of intensive studies with the goal of inducing regeneration to treat human retinopathies [16]. Although those recent studies have provided exciting new insights into the gene regulatory networks that contribute to species-specific Müller glial cell response to stress, injury, and disease, the underlying molecular mechanisms remain elusive. Moreover, it is not known how those differences are encoded during development.

Comprehensive genome-wide analyses have shown that chromatin state and transcriptional changes during murine and human retinal development are precisely coordinated to ensure that each cell type is born at the right time and in the right proportion during retinogenesis [17, 18]. Indeed, thousands of genes are upregulated or downregulated with temporal and spatial precision [17, 18]. Not only is the chromatin state at individual genes and enhancers precisely coordinated during retinal development, but higher order chromatin structure (euchromatin and heterochromatin) and nuclear organization are also cell-type specific [18]. Here, we analyzed the nuclear organization (euchromatin/heterochromatin), gene expression (scRNA-seq), and chromatin accessibility (scATAC-seq) of the 5 most abundant retinal cell types in murine retinae (rods, cones, amacrine, bipolar, and Müller glial cells). We discovered that the size, shape, and distribution of euchromatin and heterochromatin in the nucleus is specified by not only cell type but also chromatin looping, chromatin accessibility, and gene expression. As expected, nearly all active promoters (RNA-seq) were accompanied by open chromatin (scATAC-seq), and inactive promoters were sequestered into inaccessible chromatin domains. However, we were surprised to discover a group of genes in Müller glia that were not expressed in the normal healthy retina but had open chromatin at their promoters. These pliancy genes were epigenetically repressed in retinal neurons. Here we show that in murine and human retinae, the pliancy genes are quickly induced during stress, injury, or disease (15 conditions) and contribute to inflammation. Many of the pliancy genes are expressed early in development in retinal progenitor cells and are maintained in an open and accessible state in mature Müller glia even though they are not normally expressed in the healthy adult retina of humans or mice. By integrating our results with those of prior studies on zebrafish [16], we found that most genes required for retinal regeneration of the mammalian retina are pliancy genes in Müller glia. These results have important implications for efforts to induce Müller glial cell regeneration as a therapeutic approach to treat human retinopathies and more broadly for understanding the cellular response to stress, injury, or disease in the human retina.

## RESULTS

### Cell type–specific nuclear morphology and organization

The retina is organized into three distinct cellular layers with the rods and cones in the outer nuclear layer (ONL); the bipolar, amacrine, horizontal, and Müller glial cells in the inner nuclear layer (INL); and ganglion and displaced amacrine cells in the ganglion cell layer (GCL). The nuclei of each cell type have distinct morphological features including size, shape, and organization of euchromatin, facultative, and constitutive heterochromatin. Previous studies have shown that developmentally regulated genes can transition between the euchromatic and facultative heterochromatic domains, suggesting that nuclear morphology and organization may be developmental stage– and cell type–specific in the retina [18]. To develop a cell type–specific classifier of retinal nuclei based on their unique size, shape, and chromatin organization, we performed lattice light sheet (LLS) microscopy [19] on live cells from adult murine retina. We used 5 different GFP transgenic lines to identify rods (*Nrl-GFP*), cones *(ChrnB4-GFP)*, bipolar neurons (*Grm6-GFP*), amacrine neurons (*Gad1-GFP*), and Müller glia (*Rlbp-GFP*) (Fig. 1A-E) [20–24]. The DNA was stained with SiR-DNA (Cytoskeleton, Inc.), and cells were imaged using a 7.5-mm light sheet with a hexagonal pattern in sample scanning and dithered acquisition modes (Supplemental Information). To quantify the euchromatin and the facultative and constitutive heterochromatin domains across cell types, we utilized supervised segmentation and machine learning in the ilastik software pixel classification package (Supplemental Information). To ensure consistent segmentation across the entire data set, images were cropped to include only single cells and normalized in Fiji to standardize the intensity of chromatin labeling (Supplemental Information). Regions of high, medium, and low chromatin density were manually annotated in a subset of cells for each cell type to serve as a training dataset in ilastik. The resulting 3D segmentation classifier was then applied to the entire data set. In total, 234 nuclei were analyzed across 10 experiments (Table S1). As expected, the rods had the smallest nuclear volume (66±16 μm^3^) and the amacrine cells had the largest (163±44 μm^3^) (Table S1). Similarly, rods had the smallest volume and proportion of euchromatin (14±10% euchromatin) and amacrine cells had the largest (48±9% euchromatin) (Table S1, Figure S1).

**Figure 1.**
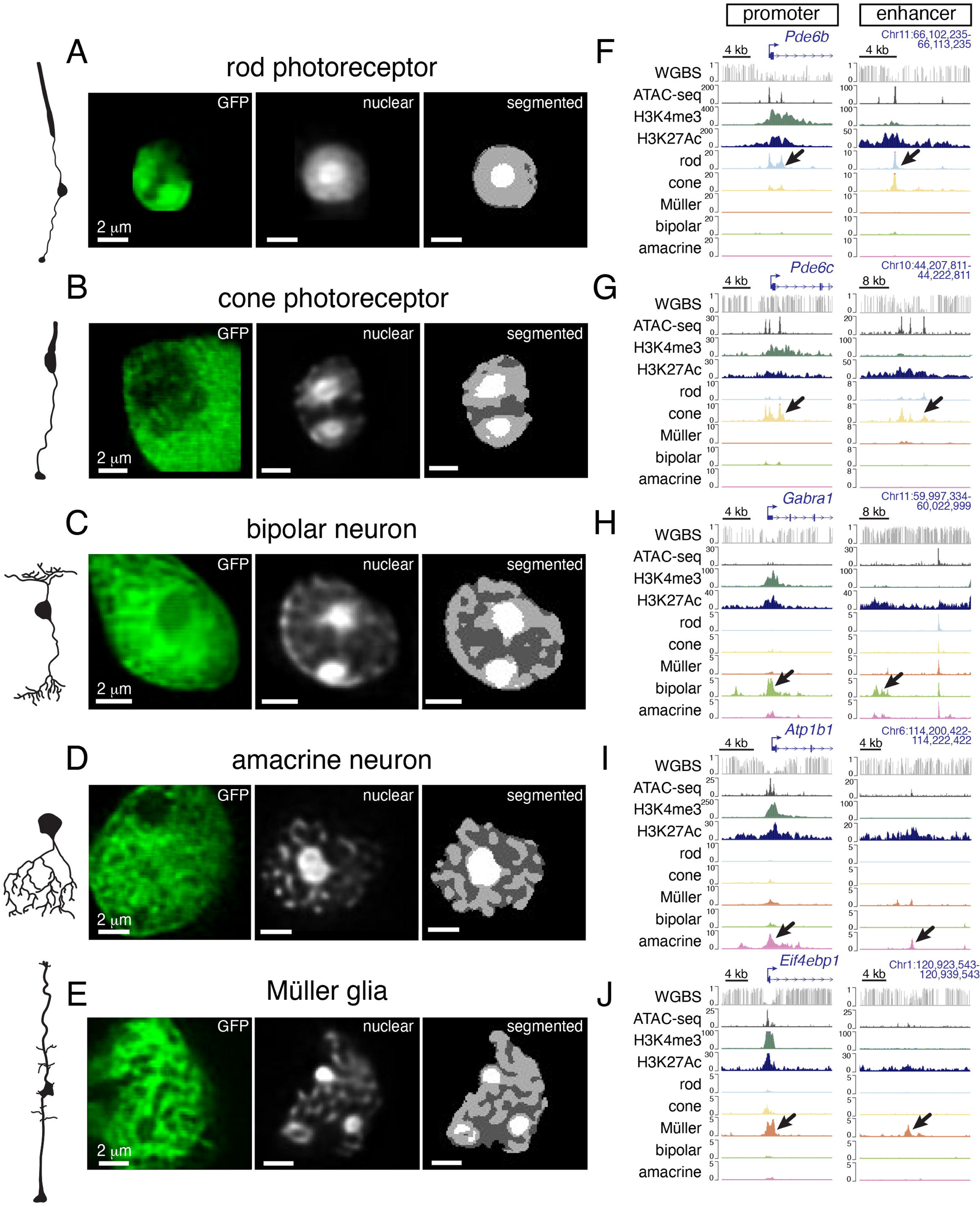
Retinal neurons and glia have cell type–specific nuclear structure and chromatin accessibility. **A-D)** Micrographs of live dissociated retinal cells expressing GFP (green) and stained with SiR-DNA to label the nuclear chromatin. Automated segmentation of euchromatin (dark gray), facultative heterochromatin (light gray), and constitutive heterochromatin (white) is shown for each nucleus. **F-J)** Tracks for whole-genome bisulfite sequencing (WGBS), bulk ATAC-seq, bulk ChIP-seq, and single-cell ATAC-seq (rod, cone, Müller, bipolar, amacrine) for representative promoters and enhancers that have cell type–specific chromatin accessibility (arrows) that correlates with cell type–specific expression (promoters) or activity (enhancers). The scale is shown for each track. Abbreviations: kb, kilobase; GFP, green fluorescent protein.

Beyond the differences in total volume and proportion of euchromatin/heterochromatin, the organization of subnuclear domains is also cell-type specific. For example, rods have a single centrally located domain of constitutive heterochromatin, and Müller glia have 2 or 3 distinct constitutive heterochromatin domains on the nuclear periphery (Fig. 1A-E). To determine whether the size, shape, and organization of nuclear domains are sufficient to distinguish retinal cell types, we used a subset of the data to train a neural network and then tested the network’s accuracy on the remaining data (Supplemental Information). Specifically, the data were used to train a random forest machine learning algorithm, which used 67% random selection of images as a training set and the remaining 33% for validation. We repeated training and validation of the model for 2,000 iterations and summarized the performance using a confusion matrix for each of the 234 nuclei in the dataset (Figure S1). Rod nuclei were identified with 100% accuracy due to their unique nuclear size, shape, and organization. Cone, bipolar, amacrine, and Müller nuclei were correctly identified 78%, 72%, 58%, and 73% of the time, respectively. The accuracy improved to 88% (14/16 cone), 77% (10/13 amacrine), 94% (32/34 bipolar) and 88% (30/34 Müller) when we used a minimum threshold of 60% confidence (Supplemental Information). Expectedly, the amacrine cells have the lowest accuracy because they have the greatest subtype diversity [25]. Taken together, these data suggest that nuclear size, shape, and organization of chromatin is cell-type specific in the murine retina.

### Cell type–specific chromatin accessibility and gene expression

To relate the cell type–specific nuclear morphology to individual genes, enhancers, and chromatin domains, we utilized our integrated epigenetic dataset (whole genome bisulfite sequence, RNA-seq, ATAC-seq, ChIP-seq, Hi-C, ChromHMM) [17, 18]. Here, we add scRNA-seq and scATAC-seq (10X Genomics) to provide cell type–specific resolution and Hi-ChIP for promoter-enhancer interactions. In total, data from more than 2,500 files are freely shared on the St. Jude Cloud in an interactive viewer (https://proteinpaint.stjude.org/iRNDb_v2). Cell type– specific chromatin accessible regions were identified at promoters and enhancers (Fig. 1F-J). For example, *Aipl1* is not expressed during embryonic stages but is rapidly upregulated in differentiating rods and cones. This pattern is reflected in the bulk retina chromatin profiling results, and the scATAC-seq and scRNA-seq results provide insights into the cell type in which *Aipl1* is upregulated (Fig. S2A-C). In contrast, the *Sfrp1* gene is highly expressed in retinal progenitor cells in embryonic retina with associated active chromatin marks but is rapidly downregulated as cells differentiate during development (Fig. S2D-F). The scATAC-seq and scRNA-seq demonstrates that *Sfrp1* is expressed in retinal progenitor cells and newly postmitotic cells (neurogenic) but is not expressed in any early-born cell types such as cones (Fig. S2D-F). Moreover, the *Sfrp1* promoter is repressed in the adult retina with H3K27me3 modifications, and this leads to reduced chromatin accessibility across all retinal cell types in the mature retina (Fig. S2E,F). These examples highlight the importance of integrating single-cell profiling with the bulk analyses.

The majority (6,614/7,083; 93%) of genes that were expressed in individual cell types (see Fig. S2) or across multiple cell types (Fig. 2A,B) also had ATAC-seq peaks at the corresponding promoter in the same cell type (Table S2, Fig. S3). The remaining 469 genes that were expressed in each cell type but lacked an scATAC-seq peak at the promoter in the same cell type were below the threshold set for our analysis but had matched expression and ATAC-seq for cells with the highest expression (Table S2). Surprisingly, a group of 364 poised genes had cell type–specific scATAC-peaks at their promoter but no expression in scRNA-seq or bulk RNA-seq (FPKM<1.0) (Table S3). For example, the *Pdk4* promoter is polycomb repressed (H3K27me3) and the gene is not expressed at any stage of retinal development (Fig. 2C). However, the scATAC-seq data revealed that the chromatin at the promoter is open and accessible in Müller glia (Fig. 2D). Most of the genes with the unusual chromatin-accessible but transcriptionally silent (accessible/silent) promoters (304/364; 83%) were in Müller glia (Fig. 2E,F). Enrichr [26] results indicated that the most significant pathways associated with the 304 genes in Müller glial accessible/silent promoters were related to inflammation (TNFα, NF-κB), angiogenesis (VEGF, HIF1α), and signal transduction (WNT/β-catenin signaling).

**Figure 2.**
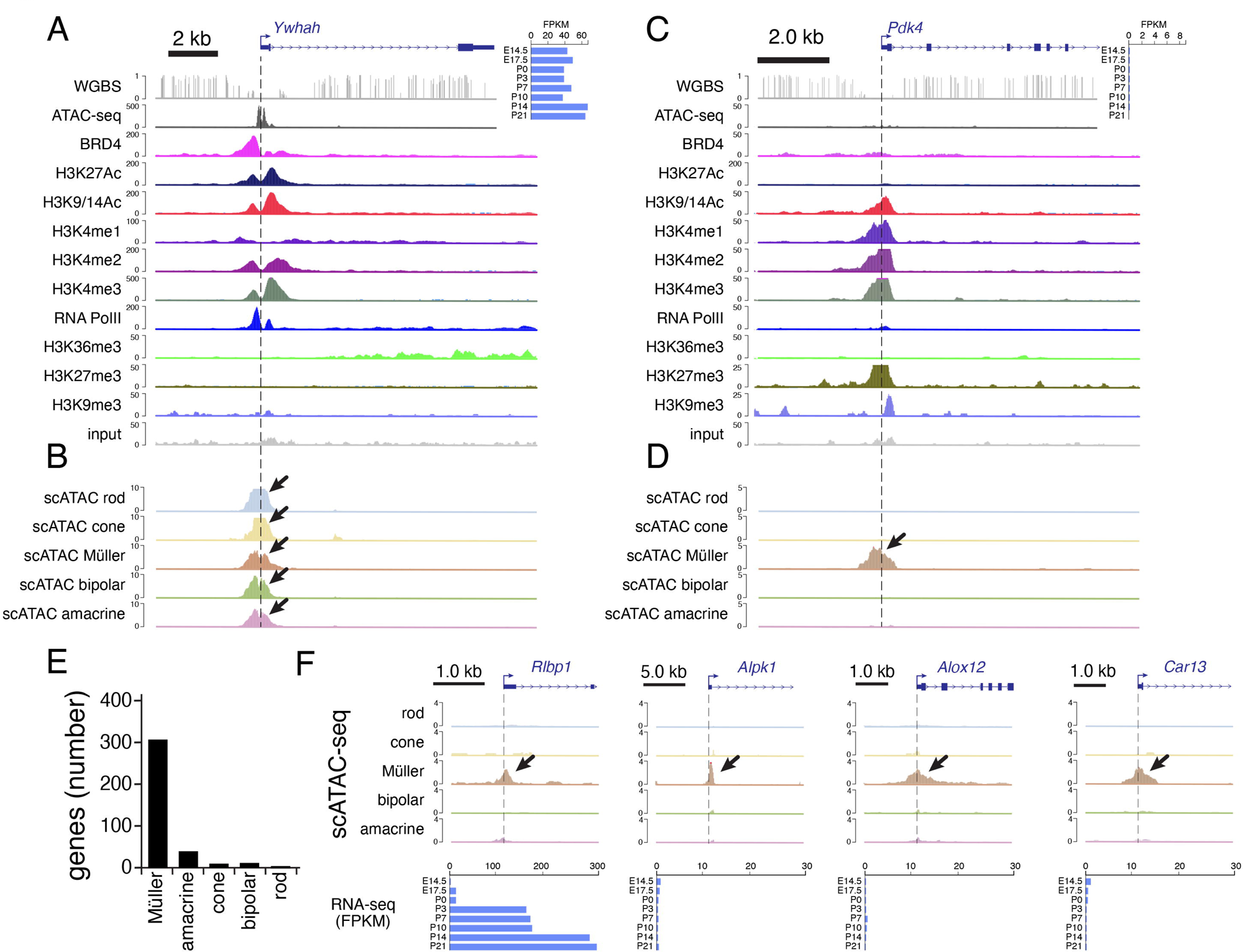
Müller glia have genes with accessible promoter chromatin that are transcriptionally silent. **A)** Tracks for DNA methylation (WGBS), ATAC-seq, and ChIP-seq for a representative gene that is expressed throughout development (horizontal bar plot in upper right). **B)** Single-cell ATAC-seq (scATAC-seq) showing accessible chromatin in each cell type (arrows), consistent with expression in those cells shown by single-cell RNA-seq (not shown). **C)** Tracks for DNA methylation (WGBS), ATAC-seq, and ChIP-seq for a representative gene that is not expressed at any stage of retinal development (horizontal bar plot in upper right). **D)** scATAC-seq showing accessible chromatin in Müller glia (arrow). **E)** Bagplot showing the number of genes with cell type–specific accessible chromatin (scATAC-seq peaks) but lacking expression in each retinal cell type. **F)** Müller-specific scATAC-seq for 4 genes: 1 gene that is expressed in Müller glia (*Rlbp1*) and 3 that are accessible/silent (*Alpk1, Alox12, Car13*). The horizontal bar plot below each set of tracks shows the expression across development. Abbreviations: FPKM, fragments per kilobase per million reads; kb, kilobase.

### Cell type–specific response to stress, injury, and disease in murine and human retinae

To determine whether any of the genes with accessible/silent promoters in Müller glia are acutely activated during stress, injury, or disease, we developed a workflow using human and murine retinal tissue punches (Supplemental Information). Briefly, 1.5-mm retinal punches were placed on polycarbonate filters in explant culture medium as described previously [14, 27–32] (Supplemental Information) (Fig. 3A). The explant cultures were then exposed to 15 different conditions to mimic stress, injury, or disease (Fig. 3A). For controls, we used the original uncultured tissue samples and retinal punches cultured with no perturbation. After 24 hours, we performed bulk RNA-seq on 2 or 3 biological replicates for each condition for murine (41 samples) and human retinae (46 samples) (Fig. 3A, Table S4, S5). In addition to performing these 87 bulk RNA-seq experiments, we also performed 46 single cell RNA-seq (scRNA-seq) experiments (29 murine, 17 human) with a total of 172,758 cells (90,730 murine, 82,028 human) that passed quality control (Table S6).

**Figure 3.**
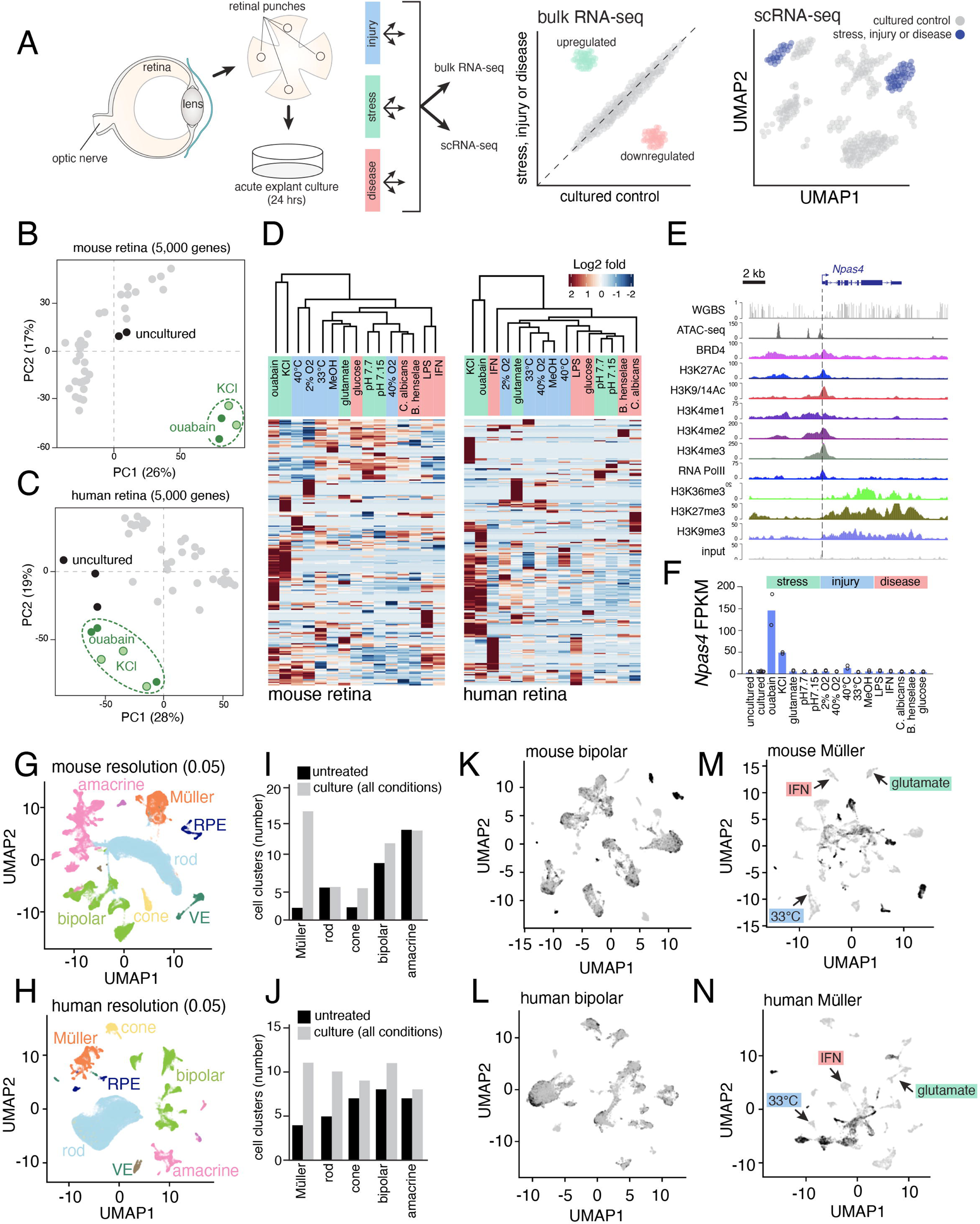
Müller glial cell–specific response to stress, injury, and disease in human and murine retinae. **A)** Drawing of the experimental workflow. Duplicate retinal punches were explant-cultured for 24 hours under 15 different conditions that mimic ocular stress, injury, or disease. Fresh tissue and cultured retinae were used as controls. Each sample was analyzed by bulk RNA-seq and scRNA-seq. **B,C)** Principal component analysis of bulk RNA-seq for mouse and human retinae. KCl and ouabain (green) caused similar changes in gene expression that were distinct from those caused by other treatments and found in uncultured retinae. **D)** Unsupervised hierarchical clustering of the bulk RNA-seq for each condition for mouse and human retinae. **E)** Representative DNA methylation, ATAC-seq, and ChIP-seq for a gene (*Npas4*) that is specifically upregulated following ouabain or KCl exposure. **F)** Bar plot of FPKM values for each of the 15 treatment conditions for the *Npas4* gene. Bars represent the mean, and the circles represent individual samples. **G,H)** UMAP of scRNA-seq for mouse and human retinae data for all conditions. **I,J)** Individual cell populations were isolated and re-clustered to give the number of clusters on the bar plot for untreated (black) and cultured (gray) murine (I) and human (J) retinal samples (all conditions combined). **K,L)** UMAP of clustered bipolar neurons for untreated (black) and treated (gray) samples. The distinct bipolar cell clusters of treated and untreated samples are similar. **M,N)** UMAP of clustered Müller glia for untreated (black) and treated (gray) samples. Distinct clusters are identified (arrows) for different treatment conditions that are separate from the uncultured or control cultured conditions. Abbreviations: FPKM, fragments per kilobase per million reads; IFN, interferon; VE, vascular endothelial cells; RPE, retinal pigment epithelium; PC principal component; UMAP, uniform manifold approximation and projection; kb, kilobase.

For the stress group, we used 70 μM ouabain or 80 mM KCl to simulate prolonged neuronal depolarization [33–37], 1 mM glutamate to simulate excitotoxicity [38–40], and extracellular pH alterations (pH 7.7 for alkalosis and pH 7.15 for acidosis) to perturb synaptic activity [41–45]. For the injury group, retinal punches were exposed to hypoxia (2.3% O_2_) [46–48], hyperoxia (40% O_2_) [49–51], hyperthermia (40°C) [52–54], hypothermia (33°C) [55–57], and methanol poisoning (8 μM formic acid) [58–61]. In the disease category, retinae were exposed to lipopolysaccharide (LPS, 0.2 μg/mL), mimicking Gram-negative bacterial infection [62–65]; interferon-γ (1.0 μg/mL), mimicking inflammation [66–68]; *Candida albicans* (100 CFU/mL), mimicking ocular candidiasis [69–72]; *Bartonella henselae* (100 CFU/mL), mimicking neuroretinitis [73]; and hyperglycemia (20 mM glucose), mimicking diabetes [74].

Retinal response was specific to the type of perturbation and is evolutionarily conserved (Fig. 3B-D and Tables S4, S5). For example, the neuron-specific immediate early gene involved in neuroprotection (*Npas4*) was upregulated 50-to100-fold in retinae treated with ouabain or KCl (Fig. 3E,F and Tables S4, S5, S6) [75]. Across all stresses, we identified 3,272 genes in the human retina and 4,799 genes in the mouse retina that were upregulated 2-fold or more in at least one condition in the bulk RNA-seq data (Table S4, S5). In contrast, 3,089 genes in the human retina and 4,863 genes in the mouse retina were downregulated 2-fold or more in at least one condition (Tables S4, S5). We identified 3,184 mouse orthologues for the 3,272 human genes that were upregulated, and 55% (1,758/3,184) were upregulated in both species by at least 2-fold (Table S5). Similarly, we identified 2,997 mouse orthologues for the 3,089 human genes that were downregulated, and 51% (1,536/2,997) were downregulated in both species by at least 2-fold (Table S5). For the described genes with accessible/silent promoters, 73% (267/364) were upregulated 2-fold or more in at least one condition, and 86% (230/267) of those had Müller glial cell–specific accessible promoters. Enrichr identified the top ontologies as inflammation, immune regulation, and vasculature.

Next, we analyzed the scRNA-seq data from mouse and human retinae exposed to the same 15 conditions already described. In total, 90,730 murine cells and 82,028 human cells passed quality control (Table S6). As for the previous study, we compared uncultured retinae to cultured retinae to eliminate the gene expression changes that result from explant culture but are not specific to the stress response (Table S6). The cell type–specific changes in gene expression in response to stress are in Müller glia and are consistent with the bulk RNA-seq data (Fig. 3G-N and Tables S7,S8). The Müller glia had more distinct clusters of cell states based on individual treatments than all other cell types did. For example, 2 clusters of Müller glia were found in the untreated murine retinae, and this increased to 17 clusters following the 15 treatments. In contrast, the number of rod clusters remained the same (n=6) (Fig. 3G-J). Similar trends were found with human retinae (Fig. 3 K-N).

Not only did the 15 stimuli induce cell type–specific responses in Müller glia, but they were specific to the individual stimulus and category. The activity-regulated cytoskeletal-associated protein encoding gene (*Arc*) was upregulated primarily in Müller glia in response to stress (KCl) but not injury (40% O_2_) or disease stimuli (hyperglycemia) (Fig. 4A). The bulk RNA-seq data were consistent with the scRNA-seq data (Fig. 4A and Tables S4, S7). Arc is involved in activity-dependent neuronal plasticity in the brain [76] as well as acute and chronic neuroinflammation [77]. We found no prior reports of Arc function or expression in the retina; yet *Arc*’s upregulation in Müller glia following KCl treatment is consistent with its role in other regions of the CNS.

**Figure 4.**
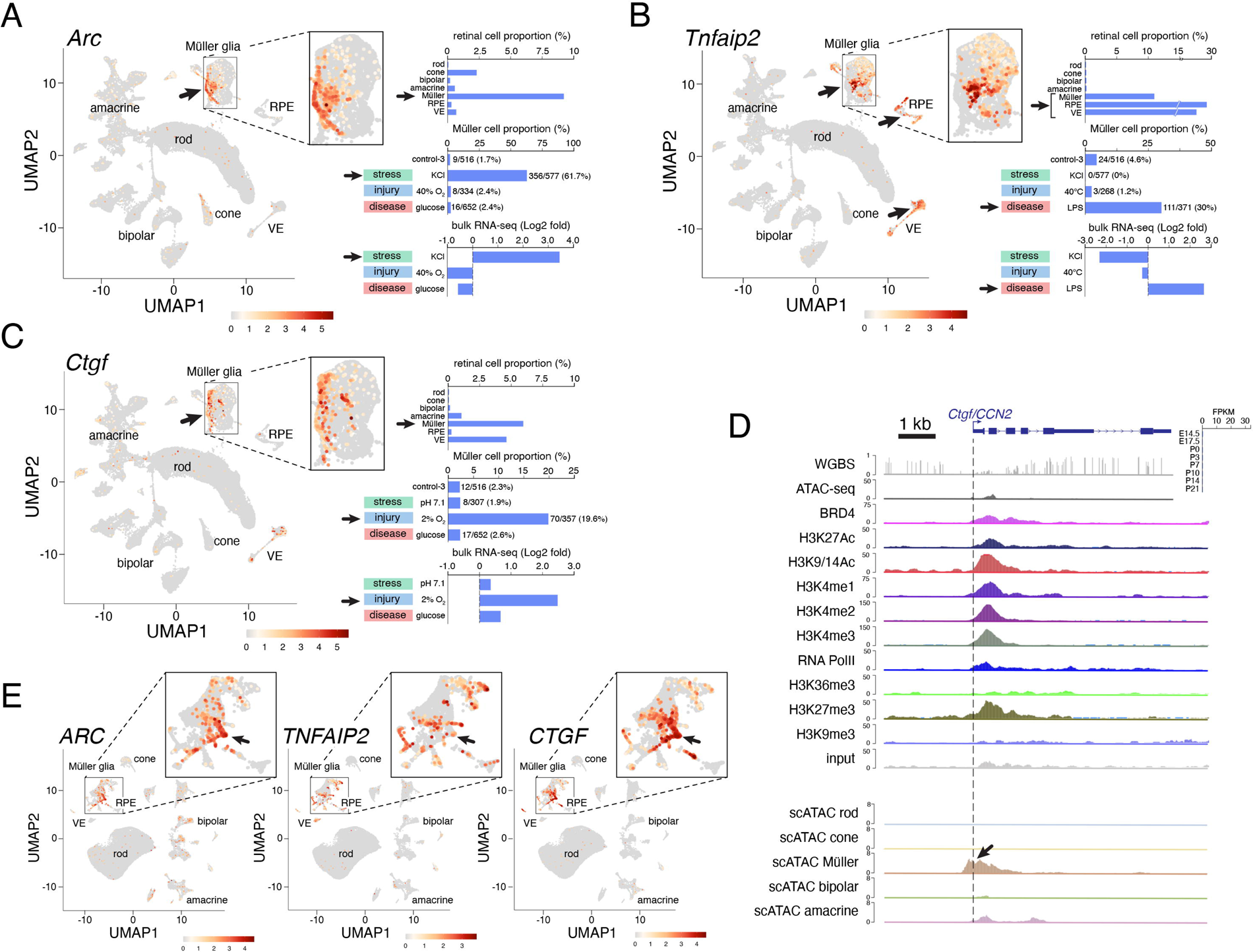
Müller glia–specific response to stress, injury, or disease conditions in human and murine retinae. **A)** *Arc* is specifically upregulated in Müller glia during retinal stress (KCl). UMAP of single-cell RNA-seq for murine retinae, including uncultured cells, control cultures, and 15 conditions mimicking stress, injury, and disease conditions showing *Arc* expression (red). Horizontal bar plots show the proportion of retinal cell types expressing *Arc* (upper), the proportion of Müller glia expressing *Arc* (middle), and the Log2-fold change relative to control cultured retinae (lower). Arrows indicate data for KCl exposure and for comparison, a representative injury (40% O_2_) and disease condition (high glucose) are shown. **B)** *Tnfaip2* is specifically upregulated in Müller glia during retinal disease (LPS). UMAP of single-cell RNA-seq for murine retina including uncultured cells, control cultures, and 15 conditions for stress, injury, and disease conditions showing *Tnfaip2* expression (red). Horizontal bar plots show the proportion of retinal cell types expressing *Tnfaip2* (upper), the proportion of Müller glia expressing *Tnfaip2* (middle), and the Log2 fold change relative to control cultured retina (lower). Arrows indicate data for LPS exposure, and for comparison, a representative injury (40% O_2_) and stress condition (KCl) are shown. **C)** *Ctgf* is specifically upregulated in Müller glia during retinal injury (2% O_2_). UMAP of single-cell RNA-seq for murine retina including uncultured, control cultures and 15 conditions mimicking stress, injury, and disease conditions showing *Ctgf* expression (red). Horizontal bar plots show the proportion of retinal cell types expressing *Ctgf* (upper), the proportion of Müller glia expressing *Ctgf* (middle), and the Log2 fold change relative to control cultured retina (lower). Arrows indicate data for LPS exposure, and for comparison, a representative disease (high glucose) and stress condition (pH7.1) are shown. **D)** Tracks for DNA methylation (WGBS), ATAC-seq, and ChIP-seq for *Ctgf*. Bagplot of expression during development from bulk RNA-seq is shown in the upper right. The lower panel shows the scATAC-seq with Müller glial cell–specific accessible chromatin at the promoter. *Ctgf* is an accessible/silent pliancy gene. **E)** UMAP for human retina for *ARC, TNFAIP2,* and *CTGF* showing evolutionary conservation of the specific patterns of expression following stress, injury, or disease conditions. Abbreviations: VE, vascular endothelial cells; RPE, retinal pigment epithelium; kb, kilobase; FPKM, fragments per kilobase per million reads.

In contrast to *Arc*, the tumor necrosis factor alpha–induced protein 2 gene (*Tnfaip2*) gene was not upregulated following KCl exposure but was upregulated following disease stimulus (LPS) (Fig. 4B). Müller glia, vascular endothelial (VE) cells, and retinal pigment epithelial (RPE) cells upregulated *Tnfaip2* following LPS exposure, and this was not observed with stress (KCl) or injury (40°C heat shock) in scRNA-seq or bulk RNA-seq analyses (Fig. 4B and Tables S4 and S7). As with Arc, Tnfaip2 expression or function has not been described in the retina.

Tnfaip2 is involved in inflammation, angiogenesis, cell proliferation, and migration [78]. In addition to finding these examples of stress stimuli (KCl) and disease stimuli (LPS), we also identified Müller-specific responses to injury. The connective tissue growth factor gene (*Ctgf/Ccn2)* has an accessible/silent promoter and is upregulated under hypoxic stimuli (2% O_2_) but not under stress (pH 7.1) or disease (hyperglycemia) (Fig. 4C,D and Tables S4 and S7). The Ctgf/CCN2 protein is a secreted growth factor that regulates multiple signaling pathways involved in cell adhesion, migration, angiogenesis, and matrix reorganization during disease processes [79]. Ctgf/CCN2 plays a role in reorganization of retinal vasculature in rd10-degenerating murine retinae [80]. These patterns of cell type–specific and damage-specific response are conserved in the human retinae (Fig. 4E and Tables S5 and S8). Hereafter, we refer to these Müller glial cell–specific genes with accessible/silent promoters as “pliancy” genes because they contribute to the damage-specific response of Müller glia to stress, injury, or disease.

### Müller glial cell pliancy genes contribute to inflammation in vivo

Müller glia regulate retinal homeostasis and contribute to retinal inflammation through cytokine/chemokine signaling and interactions with retinal vasculature and microglia [81]. Their proximity to the retinal vasculature and the blood retinal barrier make them an ideal conduit for signaling between the retina and the immune system during stress, injury, and disease to modulate inflammation [81]. Consistent with this role, we found that the most prevalent pathways for the pliancy genes that were upregulated at least 2-fold in Müller glia across all stimuli were inflammation (52%, 70/135 genes), vasculature (30%, 40/135) and NFκB/IFNγ/TNFα signaling (11%, 15/135) (Table S9). Therefore, to extend our studies in vivo, we used the well-established model of endotoxin-induced uveitis (EIU) that involves inducing ocular inflammation with an intravitreal injection of lipopolysaccharide (LPS) [82]. This not only allows us to directly compare the results from in vivo to ex vivo LPS exposure but also is clinically relevant as uveitis contributes to 5-10% of vision loss worldwide [83]. Previous studies have demonstrated that inflammation peaks 24 hours after intravitreal injection of LPS [84] and is accompanied by increased chemokine/cytokine production [85–87], increase vascular permeability [88, 89], immune cell infiltration [84, 90, 91] and reduction in retinal function measured by electroretinogram (ERG) [92–94]. For example, previous studies have shown that *Nr0b2 (Shp1), Il1b, Ccl2, Ccl3, Cxcl1*, and *Il6* are upregulated in ocular inflammation [86, 93], and each of those genes are upregulated following one or more of the stress, injury, or disease stimuli in our study (Tables S4, S5).

For EIU, 1 μL LPS (25 ng/μL) was injected into the vitreous of C57Bl/6 mice, and 1 μL PBS was injected into the contralateral eye as a negative control. After 24 hours, we harvested the retinae and vitreous (Fig. 5A). The vitreous was mixed with fluorescent beads (AccuCount 7.0-7.9 μm, Spherotech, Inc.) for normalization of infiltrating immune cells, and the retinae were harvested to analyze changes in gene expression. Immune infiltration in the LPS-injected eyes was significantly higher than that in PBS-injected eyes, with the difference being similar in magnitude to that in previous publications on EIU (Fig. 5B) [93]. There was also upregulation of *Il6, Ccl5* and *Cxcl2* measured by qRT-PCR in the eye injected with LPS relative to the PBS injected eye (Fig. 5C). For an initial assessment of the cell populations that are infiltrating after injection of LPS, we performed scRNA-seq and used the automated annotation of cell populations in SingleR [95]. We found that 42% of the infiltrating cells were granulocytes; 41%, monocytes; 6%, B-cells; and 11%, T/NK-cells (Fig. 5D). To further refine the cell populations, we performed flow cytometry (Supplemental Information), with spleen as a positive control and PBS-injected vitreous samples as a negative control (Fig. 5E). Consistent with prior studies, the proportion of granulocytes (neutrophils) among all CD45+ cells were increased in two biological replicates (Fig. 5E) [91, 96]. Monocytes were also increased in one of the replicates (Fig. 5E). However, the underlying molecular mechanisms or cellular source of signaling to recruit neutrophils during uveitis has not been elucidated.

**Figure 5.**
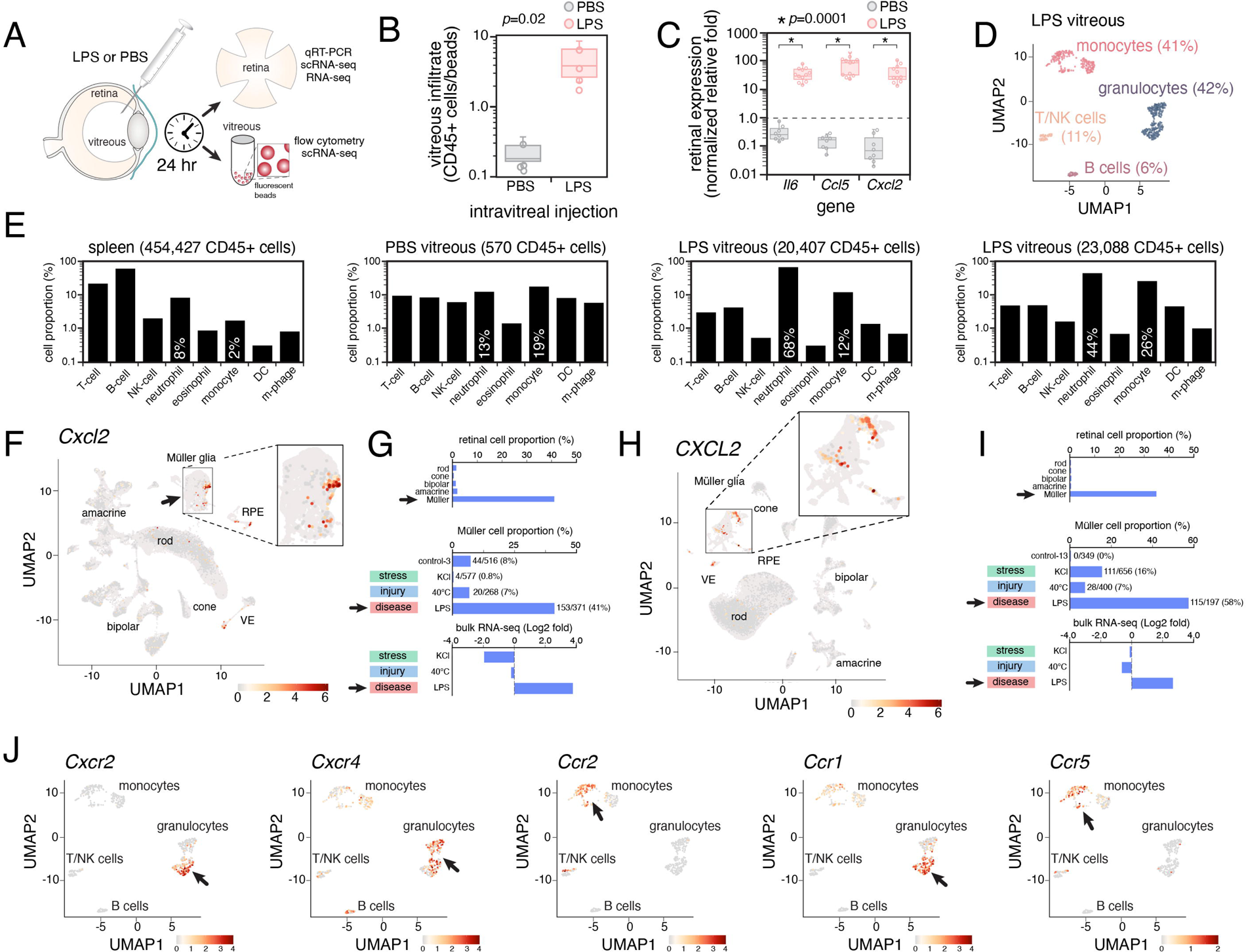
Müller glia regulate inflammation in vivo by activating pliancy genes. **A)** Drawing of experimental workflow. LPS is injected into the vitreous of the eye; 24 hours later, the retina and vitreous are harvested separately. The retinae are used for bulk and single-cell RNA-seq, and the vitreous is mixed with fluorescent beads and used for flow cytometry to measure immune cell infiltration. **B)** Box and whisker plot of the number of CD45+ immune cells per fluorescent bead following LPS or PBS injection. Individual eyes are represented by circles. **C)** Box and whisker plot of gene expression measured by qRT-PCR of retinae following LPS or PBS injection. Data are normalized to *Gapdh* expression relative to untreated retinae. Individual eyes are shown with circles, and these correspond to the same samples as in (B). **D)** UMAP of scRNA-seq for immune infiltrate cells in the vitreous 24 hours after LPS injection. **E)** Bar plots showing the proportion of CD45+ cells corresponding to each immune cell population. Spleen is used as a positive control for each cell population, and PBS is a negative control for the eye injection. Two representative eyes are shown for LPS. The total number of CD45+ cells is shown at the top of each plot. There are very few CD45+ cells in the PBS-injected eye because immune cells are not normally present in the eye. **F)** *Cxcl2* is specifically upregulated in Müller glia during retinal disease (LPS). UMAP of single-cell RNA-seq for murine retinae including uncultured cells, control cultures, and 15 conditions mimicking stress, injury and disease conditions showing *Cxcl2* expression (red). **G)** Horizontal bar plots showing the proportion of retinal cell types expressing *Cxcl2* (upper), the proportion of Müller glia expressing *Cxcl2* (middle), and the Log2 fold change relative to control cultured retina (lower). Arrows indicate data for LPS exposure, and for comparison, a representative stress (KCl) and injury (40% O_2_) are shown. **H,I)** The data for *CXCL2* in human retina are shown. **J)** UMAP for expression of chemokine receptors in the immune cell infiltrate following LPS injection. Most infiltrating cells are granulocytes and monocytes, and they express the receptor for Cxcl2 and the other chemokines that are secreted by Müller glia. Abbreviations: LPS, lipopolysaccharide; PBS, phosphate buffered saline; UMAP, uniform manifold approximation and projection; VE, vascular endothelial cells; RPE, retinal pigment epithelium; NK, natural killer cells; DC, dendritic cells.

Chemokines play an essential role in modulating leukocyte migration and invasion in a tissue-specific manner based on the stress, injury, or disease [97, 98]. Following LPS exposure in human and murine retinae, a specific subset (*Cxcl2, Cxcl12, Cxcl16, Ccl2, Ccl5, Ccl7*) of the 30 murine chemokine genes was upregulated (Table S10) in the Müller glia, and neutrophils/monocytes were recruited to the retina and vitreous (Fig. 5D,E). Some of the chemokine genes are pliancy genes (Fig. S4), and the response is specific to the perturbation. For example, murine and human *Cxcl2/CXCL2* are upregulated in Müller glia following LPS exposure (EIU) as measured by scRNA-seq and bulk RNA-seq but were not upregulated under conditions of stress (KCl-induced prolonged neuronal depolarization) or injury (40°C heat shock) (Fig. 5F-I). These gene expression data were verified by using a membrane cytokine immune assay (Supplemental Information and Fig. S5). These chemokines are relevant because their receptors (*Cxcr2, Cxcr4, Ccr1, Ccr2, Ccr5*) are expressed on the granulocytes and monocytes that infiltrate the vitreous after in vivo LPS injection (Fig. 5J).

### Cxcl2 from Müller glia signals through Cxcr2 on neutrophils during inflammation

Neutrophils are the principal cellular arm of innate immunity and one of the first lines of defense during inflammation[99]. Directed neutrophil migration following stress, injury, or disease requires 3 steps. The neutrophils must first be captured and adhere to the vascular endothelial cells near the site of injury. Next, they undergo trans-endothelial migration, passing through the basement membrane and pericytes. Finally, the neutrophils undergo interstitial motility within the tissue. Chemokines and their receptors play a critical role in tissue-specific neutrophil recruitment during inflammation and can signal through both paracrine and autocrine mechanisms [100].

We have shown that Müller glia are the major source of chemokines following stress, injury, and disease in the retina. To further refine which chemokines are important for neutrophil recruitment in the EIU model, we administered blocking antibodies (α-Ccl2, α-Cxcl1, α-Cxcl2, α-Ccl5, α-Ccl7, and the pan-Cxcl antibody α-Cxcl) to the vitreous (Fig. 6A). As expected, the mRNA expression of chemokines from Müller glia following LPS administration was not affected by the blocking antibodies (Fig. 6C-D). However, immune infiltration was significantly reduced (p=0.04) following administration of the α-Cxcl2 antibody in the EIU model (Fig. 6E). To extend these results, we performed a similar experiment in mice lacking the Cxcl2 receptor (*Cxcr2*). The mRNA expression of chemokines from Müller glia following LPS administration in *Cxcr2+/+* and *Cxcr2–/–* mice was indistinguishable (Fig. 6F-H), but immune infiltration in the *Cxcr2–/–* mice was significantly less (p=0.007) than that in wild-type mice (Fig. 6I). To distinguish between autocrine and paracrine signaling, we performed bone marrow transplantation (Fig. 6J). *Cxcr2+/+* bone marrow was transplanted into irradiated *Cxcr2–/–* recipients and vice versa. All transplants were performed using male bone marrow into female bone marrow to monitor the degree of engraftment, and *Cxcr2+/+* into *Cxcr2+/+* was used as a control (Fig. 6J) (Supplemental Information). Neutrophil infiltration in *Cxcr2+/+* wild-type recipients of *Cxcr2–/–* bone marrow was significantly reduced following LPS injection and *Cxcr2–/–* recipients of *Cxcr2+/+* bone marrow restored the neutrophil infiltration following LPS administration (Fig. 6L). Taken together, these data suggest that Müller glia release Cxcl2 following LPS injection, leading to inflammation mediated by signaling through Cxcr2 on neutrophils.

**Figure 6.**
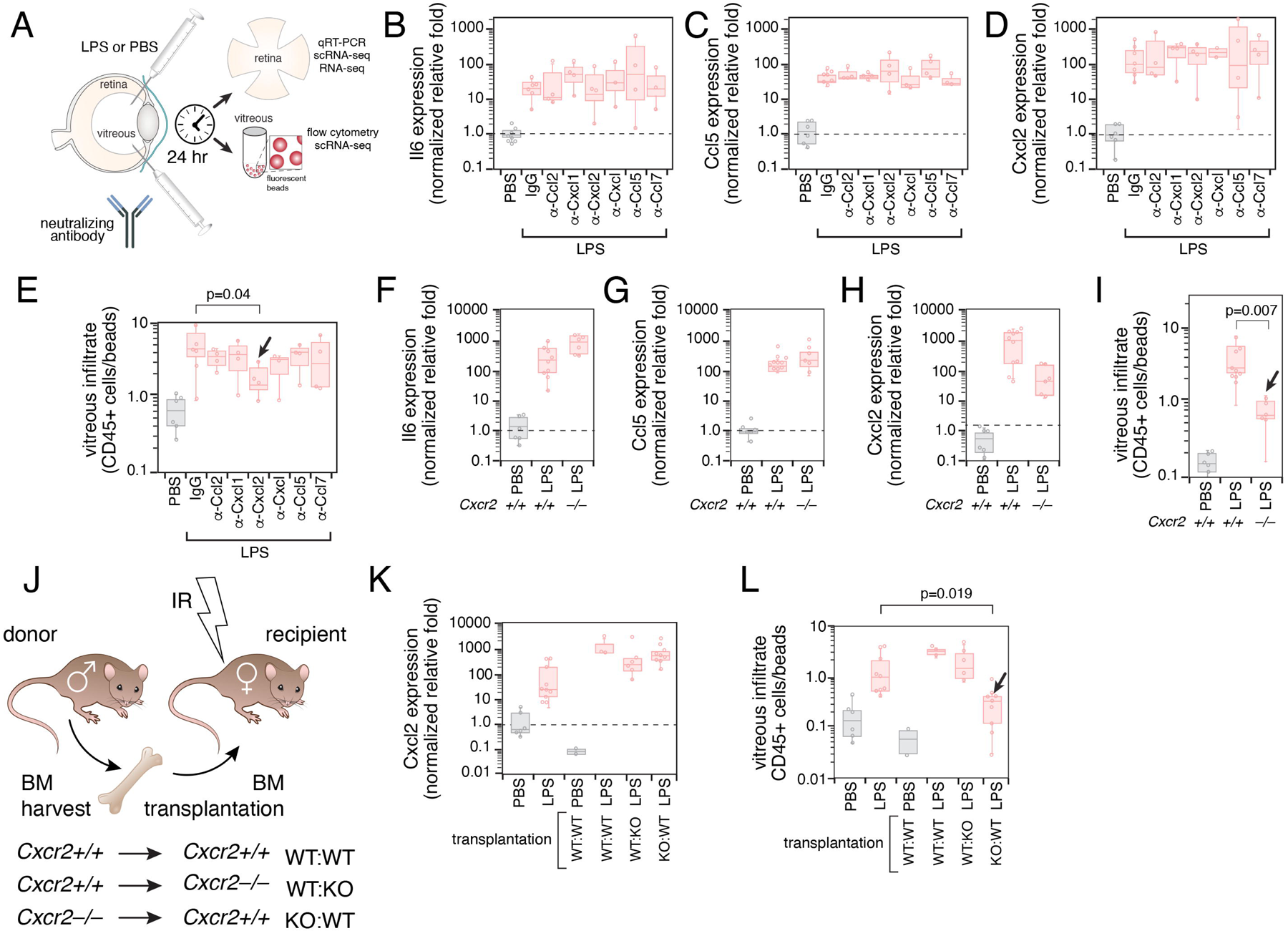
Paracrine signaling between Müller glia and neutrophils during inflammation. **A)** Drawing of the experimental workflow for testing neutralizing antibody to prevent the immune infiltration in the EIU model. In addition to LPS injection, we also injected neutralizing antibodies for individual chemokines in separate experiments, including IgG (control), α-Ccl2, α-Cxcl1, α-Cxcl2, pan anti-Cxcl (α-Cxcl), α-Ccl5, and α-Ccl7. **B-D)** Box and whisker plot with individual data points shown for qRT-PCR of Il6, Ccl5, and Cxcl2 in retinae following administration of LPS and neutralizing antibody. PBS injection is used as a control, and data are normalized relative fold. **E)** Box and whisker plot of immune cell infiltrate measure as the proportion of CD45+ cells from the vitreous after 24 hours relative to the number of fluorescent beads added to each sample. Only administration of anti-Cxcl2 neutralizing antibody led to a statistically significant reduction in immune infiltration following LPS injection (arrow). **F-H)** Box and whisker plot with individual data points shown for qRT-PCR of Il6, Ccl5, and Cxcl2 in retinae from wild-type and *Cxcr2–/–* mice following administration of LPS or PBS as a negative control. **I)** Box and whisker plot of immune cell infiltrate measured as the proportion of CD45+ cells from the vitreous 24 hours after LPS injection relative to the number of fluorescent beads added to each sample in wild-type and *Cxcr2–/–* eyes. There was a significant reduction in immune infiltration (arrow) in the *Cxcr2–/–* eyes after LPS injection relative to wild-type mice. **J)** Drawing of bone marrow transplantation experimental workflow. Male mice served as donors into female irradiated (9 Gy) mice. **K)** Box and whisker plot with individual data points shown for qRT-PCR for Cxcl2 in retinae from transplanted mice following administration of LPS or PBS as a negative control. **L)** Box and whisker plot of immune cell infiltrate measured as the proportion of CD45+ cells from the vitreous 24 hours after LPS injection relative to the number of fluorescent beads added to each sample in control and transplanted mice. There was a significant reduction in immune infiltration (arrow) in the wild-type recipients that received a *Cxcr2^−/−^*-deficient bone marrow transplantation (arrow). Abbreviations: qRT-PCR, quantitative real-time polymerase chain reaction; PBS, phosphate-buffered saline; BM, bone marrow; LPS, lipopolysaccharide.

### Comparison of human and murine Müller glia stress response to that of zebrafish

In zebrafish, the Müller glia can dedifferentiate into retinal progenitor cells and regenerate retinal neurons [101]. Therefore, we compared all of the differentially expressed genes in our bulk RNA-seq and scRNA-seq data to those identified as differentially expressed in a prior study comparing the Müller glial cell response to light damage or NMDA exposure in zebrafish and mouse retinae [16]. There was strong concordance between our dataset and that of the prior study by Hoang et al. [16]. For example, among the 354 genes that were differentially expressed (DE) in both species in Hoang et al., 38% (133/354) were upregulated by at least 2-fold in one or more conditions, and 99% (353/354) had some degree of upregulation in our experiments (Tables S4, S11). Our data extend those from the previous study by incorporating more conditions and by including human retina. Among our evolutionarily conserved upregulated genes (mouse and human), 189 were not found in the prior study (Table S11) [16]. For example, *Trim30c* (41-fold)*, Slfn4* (48-fold), and *Gbp10* (37-fold) were specifically upregulated following LPS treatment; *Chtf8* (10-fold), *Cdsn* (11-fold), and *B4galnt2* (11-fold) were upregulated at pH 7.7; and *Ephx3* (11-fold), *Utf1* (39-fold), and *Fermt3* (10-fold) were upregulated at 40°C but none of these underwent expression changes after NMDA or light damage in the prior study [16].

Several of the chemokines and inflammatory signaling genes (*Ccl2, Ccl7, Cxcl1, Cxcl10, Cxcl2, Icam1*) were found in our mouse/human data and the Hoang et al. data but lacked orthologues in zebrafish (Tables S7 and S11). However, two chemokines (cxcl18b and cxcl12b) are differentially expressed in the zebrafish retina after injury [16]. Chemokines are found in two major chromosomal locations in gene clusters (CXC and CC); although some are conserved across species, species-specific differences exist due to rapid evolution of the chemokine gene clusters [102, 103]. Chemokine orthologues can have different tissue-specific functions across species because of pathogen-driven evolutionary interactions unique to each species [102, 103]. The cxcl18b chemokine can play different roles in zebrafish depending on the condition. It is secreted by non-phagocytic cells of the stroma in response to mycobacterial infection and signals through cxcr2 to mediate pathogen-driven inflammation [104]. However, in wound-induced inflammation, cxcl18b-cxcr2 signaling is thought to be anti-inflammatory [104]. Following light damage or NMDA treatment in zebrafish retinae, cxcl18b is upregulated (Fig. S6). The cxcl12b chemokine signals through cxcr4, is pro-inflammatory, and recruits macrophages and neutrophils [104]. Following light damage or NMDA treatment in zebrafish retinae, cxcl12b is downregulated (Fig. S6). Taken together, the upregulation of the anti-inflammatory chemokine cxcl18b and the downregulation of the pro-inflammatory chemokine cxcl12b may lead to an anti-inflammatory response to retinal damage in zebrafish, which is the opposite of what occurs in murine and human retinae.

### Epigenetic state of gene networks in Müller glial cell response to stress, injury, or disease

To elucidate the changes in chromatin state following stress, injury, or disease, we performed ChIP-seq for the 8 histone marks, CTCF, RNAPolII, and Brd4 on retinae 24 hours after exposure to LPS or NMDA. The data have been made available in our online browser (https://proteinpaint.stjude.org/iRNDb_v2). Hoang et al. identified 10 distinct modules within the gene regulatory network contributing to the normal Müller glial cell states. They were further classified into resting state, reactive state after injury, return to resting state (murine), and cell-cycle re-entry/neurogenesis (zebrafish) were analyzed [16]. The genes in the murine resting-state modules (*Lhx2, Nfia*), the reactive-state modules (*Aldh1a3, Fas*), and the modules important for the return to resting (*Nfib, Sox5*) have open chromatin in Müller glia and heterochromatin silencing of the promoter in retinal neurons (Fig. 7 and Figs. S7, S8). As expected, the pliancy genes such as *Aldh1a3* that are not normally expressed but are rapidly upregulated after stress, injury, or disease are in the reactive state modules (Fig. 7B) and had more promoters bound RNAPolII after LPS or NMDA exposure (Fig. 7C and data not shown). These data suggest that reactive module gene promoters may be maintained in an open chromatin configuration to facilitate rapid changes in expression mediated by transcriptional activators and repressors (Fig. 7D,E).

**Figure 7.**
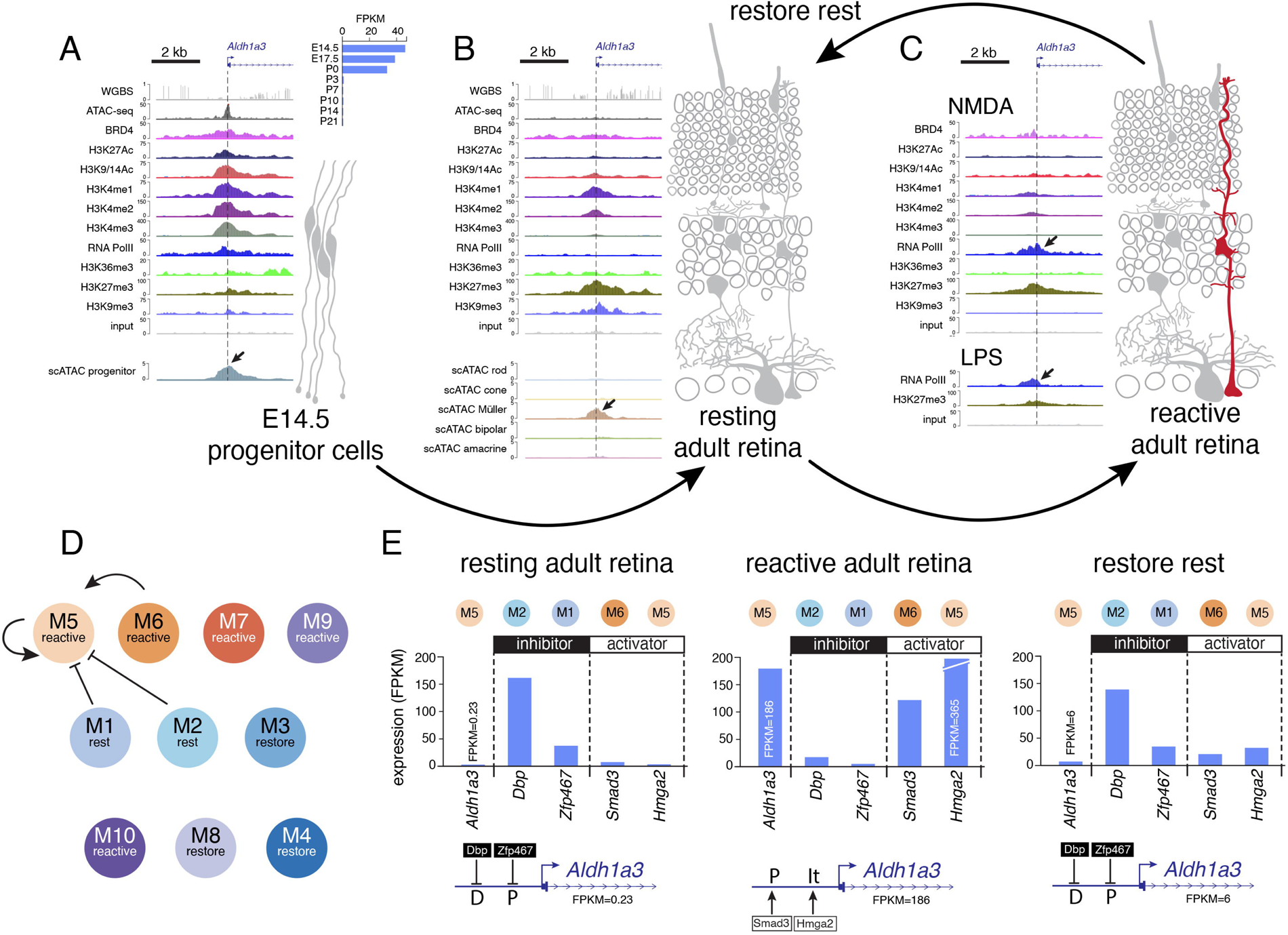
Pliancy genes are regulated by the transcriptional modules involved in Müller glia reactivity. **A)** Tracks for DNA methylation, ATAC-seq, and ChIP-seq for *Aldh1a3* gene in retinal progenitor cells at E14.5. Expression during retinal development is shown in the upper right, and scATAC-seq for retinal progenitor cells is shown in the lower panel. *Aldh1a3* is expressed in retinal progenitor cells and then silenced in the adult retina. **B)** DNA methylation, ATAC-seq, and ChIP-seq data are shown for *Aldh1a3* in the adult retina along with scATAC-seq in the lower panel. The *Aldh1a3* gene is a Müller glial cell pliancy gene. **C)** DNA methylation, ATAC-seq, and ChIP-seq data are shown for the *Aldh1a3* gene following NMDA and LPS exposure. There is an increase in RNA-PolII binding at the promoter 24 hours after NMDA or LPS exposure, consistent with an increase in gene expression. **D)** Murine Müller glial cell transcriptional modules from Hoang et al. showing *Aldh1a3* in the Müller glial cell–reactive module (M5). **E)** Expression of the *Aldh1a3* target gene and the transcription factors predicted to activate or (Smad3 (M6), Hmga2 (M5)) repress expression (Dbp (M2), Zfp467 (M1)). Abbreviations: kb, kilobase; WGBS, whole-genome bisulfite sequencing; FPKM, fragments per kilobase per million reads; LPS, lipopolysaccharide; NMDA, N-methyl-D-aspartate. P indicates promoter proximal predicted binding site; D indicates distal predicted binding site, and it indicates intragenic predicted binding site.

To determine whether the chromatin accessibility and/or chromatin landscape may contribute to the difference in activation of the proliferation/neurogenesis genes in murine versus zebrafish Müller glia, we analyzed the 270 zebrafish DE genes with mammalian orthologues in reactive state modules [16]. Surprisingly, 47% (127/270) of the zebrafish-specific reactive state module genes were upregulated in one or more of our 15 experimental conditions (Table S12). The majority (121/127) were expressed and/or had accessible chromatin at the promoter in retinal progenitor cells (Figure S9 and Table S12). Manual review of the ChIP-seq, scRNA-seq, and scATAC-seq data for all 127 genes showed two distinct patterns in the adult retina: constitutive and restricted. One group of constitutive genes (95/127) maintained the accessible chromatin pattern across retinal cell types in the adult retina and showed increased RNA-PolII binding (ChIP-seq) at the promoter after NMDA or LPS exposure (e.g., *Kif11, Atad2*) (Fig. S9 and Table S12). This group also included several cell-cycle genes (*Cdk2, Bub1, Chek1, Ccnb1, Top2a*). The other group of restricted genes (32/127) acquired heterochromatin histone modifications (H3K27me3, H3K9me3) during retinal differentiation and had Müller glia– specific accessible chromatin (e.g., *Itga2*, *Igf2bp2*) (Fig. S9 and Table S12). This group also included several neurogenic genes (*Yap1, Six3, Tgif1, Ascl1, Neurog2*) and had increased RNA-PolII binding (ChIP-seq) at the promoter after NMDA or LPS exposure (Fig. S9 and Table S12). Taken together, these data show that nearly half of the genes (127/270) that were specific for zebrafish regeneration are in an open and accessible chromatin configuration in Müller glia and can be rapidly turned on.

Next, we analyzed the TF networks from Hoang et al. [16] for the 127 genes from zebrafish regeneration modules 7 and 10 that are upregulated in our study. There are 121 transcription factors that regulate those 127 genes, but only 5 (*Sox9, Myb, Zbtb2, Rel, Atf3*) are themselves upregulated after stress, injury, or disease in our bulk RNA-seq data. We also analyzed the direct target genes and the transcription factors themselves and reviewed the scRNA-seq data. Taken together, Sox9 and Myb are predicted to regulate Ctsc, and all 3 of these genes have the restricted pattern of chromatin (see above) with robust activation in Müller glia, with Sox9 being conserved in human retina (Fig. S10).

We also analyzed the 143 genes in reactive state modules that were not turned-on in mouse retina after injury in Hoang et al. [16] or any of our stress, injury, or disease stimuli. As for the genes that are activated, the majority (137/143) were expressed and/or accessible in retinal progenitor cells (Table S12), and 135/143 had accessible chromatin in Müller glia after differentiation. Only 4 of the 270 DE zebrafish genes involved in regeneration (e.g., *Prss16*) were inaccessible and epigenetically silenced (H3K27me3, H3K9me3) in all cell types across development (Fig. S11 and Table S12). Taken together, our data suggest that the major differences between species are the transcriptional networks that control inflammation versus regeneration and that epigenetic silencing is not a major barrier to regeneration in mammalian retinae (Fig. 8).

**Figure 8.**
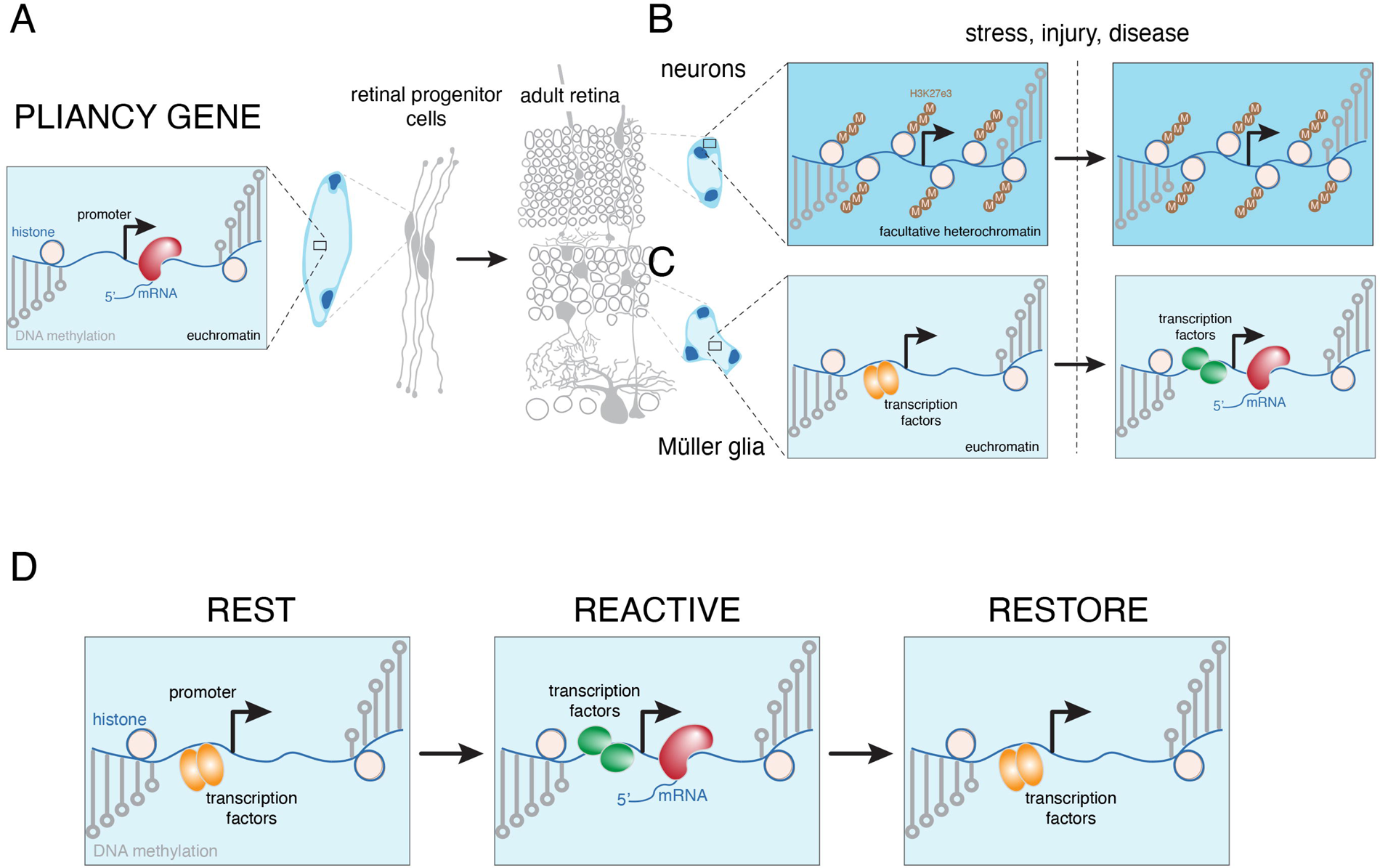
Model for cellular pliancy genes in Müller glia. **A)** Many of the cellular pliancy genes are expressed and in an open and accessible chromatin state in retinal progenitor cells. The promoters have hypomethylated DNA, histone modifications characteristic of active transcription (e.g., H3K4me3, H3K27Ac), and are bound by RNA-PolII. **B)** As retinal progenitor cells differentiate into neurons, the cellular pliancy genes are silenced by polycomb (HeK27me3) modification of histone H3. The DNA remains hypomethylated around the promoter, and the chromatin is inaccessible (ATAC-seq) and localized to facultative heterochromatin in the nucleus. In neurons, the Müller glial cellular pliancy genes are never induced in response to stress, injury, or disease. **C)** In contrast to neurons, the Müller glia maintain the promoters of the cellular pliancy genes in an open and accessible chromatin configuration and are hypomethylated. However, the gene is not expressed under normal circumstances and is only induced following stress, injury, or disease. Increased RNA-PolII binding, and transcription are hallmarks of active cellular pliancy genes in Müller glia, and these conditions are thought to be mediated by transcription factors rather than chromatin modifications. One group of transcription factors (orange) keeps the chromatin in an open and accessible state but is not sufficient to activate transcription. Following stress, injury or disease, the transcriptional circuitry changes (orange to green) and expression is activated exclusively in Müller glia. **D)** The transcriptional networks previously defined by Hoang et al. are hypothesized to be involved in retaining the pliancy gene promoters in an open and accessible chromatin state during rest. After stress, injury, or disease, the transcriptional networks are altered and lead to the reactive Müller glial cell state and gene expression. Finally, the cells return to rest by re-establishing the original transcriptional network circuitry.

## DISCUSSION

We build upon our previous epigenetic profiling studies of the developing murine retina to include scATAC-seq and scRNA-seq profiles to identify cell type–specific patterns of promoter and enhancer activity. We identified several hundred genes in Müller glia that had accessible chromatin at their promoters but lacked expression. We discovered that these accessible/silent genes were primarily in Müller glia and were upregulated in response to conditions that mimic stress, injury, or disease in human and murine retinae. We call these “pliancy genes” because they provide Müller glia with the ability to tailor their response to changes in retinal homeostasis. The functional significance of the pliancy genes in vivo was examined using the experimentally induced uveitis (EIU) model. Müller glia secrete a specific set of chemokines to directly recruit neutrophils and monocytes into the eye as part of the inflammatory response. The chemokine expression in zebrafish Müller glia is more consistent with an anti-inflammatory response. Chromatin profiling of injured murine retina and analysis of the gene regulatory network modules in zebrafish and murine retinal Müller glia suggest that the species-specific difference in response to retinal injury is regulated primarily by the transcriptional networks rather than by epigenetic silencing of the regeneration program. Taken together, our data show that during differentiation, Müller glia organize their chromatin into a pliancy program that is poised to respond to stress, injury, or disease by initiating inflammation and other injury-related pathways Transcriptional networks rather than the chromatin state contribute to the species-specific differences underlying inflammation in mammalian retinae versus regeneration in zebrafish retinae.

### Cell type–specific chromatin organization in the retina

A major limitation of our prior studies on the dynamic changes in chromatin organization during retinal development was the lack of cell type–specific information. Using lattice light sheet microscopy on live cells, we discovered that retinal cell types can be classified based on their size, shape, and organization of euchromatin and heterochromatin. The inter- and intra-cell type heterogeneity of nuclear organization contributed to the fidelity of the classifier. Rods could be confidently identified because they are homogenous and have unique features relative to the other retinal cell types, whereas amacrine cells were more difficult to classify because of the different subtypes with distinct morphologies. Additional analyses of neuronal subtypes may further refine our ability to identify retinal cell types by nuclear morphology alone. It will also be useful in future studies to determine whether groups of genes (e.g., pliancy genes) reside in proximity and/or nuclear domains.

Our cell type–specific analysis plan was to include scRNA-seq and scATAC-seq experiments. From prior study results, it was possible to identify developmentally regulated promoters and enhancers in bulk analyses, but cell type–specific profiles were lacking. We found that integrating scRNA-seq and scATAC-seq results with the bulk ChIP-seq data enabled us to predict cell type–specific chromatin patterns at promoters and enhancers. For example, some promoters have both repressive (H3K27me3) and activating (H3K27Ac, H3K4me1,2,3) histone modifications. This may represent a poised promoter or represent cell type–specific differences in chromatin organization. The scATAC-seq and scRNA-seq allowed us to better distinguish between those two possibilities. If a gene is expressed (scRNA-seq) and has accessible chromatin at the promoter (scATAC-seq) in a specific cell type but has both repressive and active histone modifications, then that is consistent with cell type–specific epigenetic silencing. In contrast, if there is no expression (scRNA-seq) or accessible chromatin at the promoter (scATAC-seq), then that is consistent with a poised promoter across retinal cell types.

The integrated data are also useful for enhancer analysis. A major challenge in the field is matching enhancers with the genes they regulate. We have been able to identify enhancers from the bulk ChIP-seq data and then identify cell type–specific co-accessible patterns to match putative enhancer-promoter interactions. Ongoing studies are focused on validating those putative promoter-enhancer interactions via Hi-ChIP and functional studies.

### Pliancy Genes in Müller Glia

We were not surprised to find that the majority (93%) of genes that were expressed in a particular cell type in scRNA-seq analyses also had accessible chromatin at their promoter in scATAC-seq analyses, but we were surprised to find genes having accessible chromatin but no expression. We confirmed the lack of expression using bulk RNA-seq, and for many of these genes, they were never expressed during retinal development. This absence suggests that there is promoter-specific chromatin organization that is not directly related to transcription. The specificity of this process was further confirmed with 3 independent observations. First, this accessible/silent pattern of promoter chromatin organization was enriched (83%) in Müller glia. Second, the genes with the accessible/silent pattern of chromatin in Müller glia were enriched in stress response pathways, including inflammation and angiogenesis. Third, these genes were rapidly induced in Müller glia in response to specific stress, injury, or disease conditions. The remarkable cell type-and damage-specific changes in gene expression led us to call these “pliancy genes” because they help the Müller glia adapt and respond to changes in retinal homeostasis that are associated with stress, injury, or disease. This pattern of accessible/silent chromatin at promoters of genes involved in response to stress, injury, or disease may have been overlooked in prior studies because of a lack of cell type–specific profiling and the focus on expressed genes. We were able to discover these genes only because of our single-cell profiling study across 15 different conditions. As mentioned, many of the pliancy genes are never expressed during retinal development, but a subset are expressed in retinal progenitor cells. For those genes, the accessible chromatin is maintained in Müller glia and repressed (H3K27me3) in the neuronal cell types. These patterns highlight the close relationship between Müller glia and retinal progenitor cells and the cell-type specificity of nuclear and chromatin organization.

### Response to Stress, Injury, and Disease

Our data on human and murine stress response extend prior studies by including 15 different types of stress, injury, or disease perturbations. We used retinal explants in short-term cultures to allow us to perform studies at scale and because some of the conditions cannot be performed in vivo. This also allowed us to use post-mortem human retinae and perform cross-species comparisons. Although the evolutionary conservation was not 100% in our experiments, the trends were consistent, and differences between the human and murine retinae were often due to the thresholds that were used to filter the data. To mitigate the effect of explant culture, we compared freshly isolated retina to control cultures. It is possible that bona-fide stress response pathways were overlooked in our study if they were also induced in the control retinal explants. The use of both bulk RNA-seq and scRNA-seq analyses was important to obtain quantitative and reproducible data (bulk RNA-seq) as well as cell type–specific data (scRNA-seq).

The genes that were previously identified [16] as being differentially expressed following NMDA exposure or light damage were also identified in our study. For example, among the 354 genes that were DE in both species by Hoang et al., 45% (159/354) were upregulated by at least Log2FC>1.0 in at least one condition in our study, and 99% (353/354) were upregulated but below the Log2FC>1.0 threshold. This was due, in part, to the comparison to cultured retina as a control, which showed some increase in expression and then mitigated the fold change.

Beyond those previously identified genes, we identified 1,385 genes that were upregulated in both mouse and human retinal explants in at least one condition but were absent from prior studies [16]. By combining the datasets, we have a broader accounting of the remarkable ability of Müller glia to respond to different conditions to maintain homeostasis. We also discovered that the neuronal response is less pronounced than that of the Müller glia. For example, among the pliancy genes in individual cell types, only 18 genes were upregulated 2-fold or more in any neuronal population in any of the 15 experimental perturbations. In contrast, 267 of the pliancy Müller glial genes were upregulated 2-fold or more after stress (211 genes), injury (150 genes), or disease (99 genes). This is important because much of the prior research in the field has been focused on the neuronal changes associated with neuronal loss in retinopathies. However, the Müller glia are also changing, and we must consider the neuronal loss in the context of the acute and sustained changes in Müller glia that are associated with the stress, injury, or disease condition.

One of the most striking results was the increased expression of a specific subset of chemokine genes in Müller glia. The major cellular process controlled by chemokines and their receptors is cell movement such as chemotaxis. In most tissues, chemokines modulate leukocyte migration and recruitment to fit the immunological needs of the tissue following stress, injury, or disease. Therefore, chemokines are central players in the inflammatory response in each tissue and under a variety of stress conditions.

Chemokine genes are found in two major chromosomal locations in gene clusters; some are conserved across species, but species-specific differences exist because of rapid evolution and species-specific adaptation to pathogens in the environment. Chemokine receptors are also organized into gene clusters, but they tend to be more evolutionarily conserved across species than the chemokine genes are. In some cases, chemokines that activate one receptor can be antagonistic to others, further increasing the complexity of chemokine function in individual tissues. Our in vivo studies with the EIU model showed a clear pattern of chemokine production by Müller glia that recruit neutrophils and monocytes. Our data are consistent with results of prior studies using the EIU model in terms of immune cell infiltration, but we show the source of the chemokines that lead to neutrophil and monocyte infiltration. We demonstrated the specificity chimeras to directly demonstrate that the Müller glia are secreting the chemokines that lead to local infiltration of immune cells. There was also evidence in our data that the vascular endothelial cells and RPE may express chemokines in the EIU model, but it was difficult to validate because of the small number of those cell types in our EIU model. Overall, these data provide a compelling link between the chromatin landscape of pliancy genes and signaling between Müller glia and the immune system following stress, injury, or disease.

### Transcriptional networks and chromatin landscape

Prior studies focused on inducing murine Müller glia to produce neurons have used pharmacologic agents that alter the chromatin landscape, such as TSA (a histone deacetylase inhibitor) [105]. The rationale for this approach is that Müller glia must undergo extensive reorganization of their chromatin architecture to de-differentiate into retinal progenitor cells and produce retinal neurons. In contrast, the more-recent studies have focused on perturbing the transcriptional modules that control the different stages of Müller glial cell response to NMDA or light damage to better recapitulate the zebrafish program in murine retina [16]. Our data from 15 different experimental conditions across murine and human retinae are more consistent with the transcriptional network model by Hoang et al. than with the chromatin reorganization model [16]. We confirm that murine Müller glia can activate proliferation (*Cdk2, Bub1, Chek1, Ccnb1, Top2a*) and neurogenic (*Yap1, Six3, Tgif1, Ascl1, Neurog2*) genes and that most of the cell proliferation genes are open chromatin without any heterochromatin (*Cdk2, Bub1, Chek1, Ccnb1, Top2a*), but the neurogenic genes (*Yap1, Six3, Tgif1, Ascl1, Neurog2*) have heterochromatin repression in neurons. These data suggest that treatment with non-specific pharmacologic agents that alter the global chromatin landscape may not be required for Müller glial cells to dedifferentiate and produce retinal neurons, but it may be counterproductive as it may harm the neurons that do survive. Indeed, all but 4 of the DE genes in zebrafish that are involved in proliferation and regeneration have promoters in an open and accessible chromatin configuration in Müller glia in murine and human retinae. Thus, future studies should continue to focus on rewiring the transcriptional modules (e.g., *Nfia/b/x*) to shift from inflammation to dedifferentiation if retinal regeneration from Müller glia in mammalian retina is to be a possibility for treatment of human retinopathies. Moreover, these modules should also be the focus of efforts to modulate the ocular inflammatory response and neovascularization because we have shown here that the Müller glia are the primary source of signals.

## ACKNOWLEDGEMENTS

We thank the MidSouth Eyebank for providing human retinae. We thank Wendy Lin, Thomas Reynoldson, Natalie Geiger, and Everest Ouyang for technical assistance. We thank the Flow Cytometry Shared Resource (Scott Perry) for assistance with immune infiltration analysis and the Comparative Pathology Shared Resource (Heather Sheppard) for analysis of bone marrow samples. We thank Cherise Guess for editing the manuscript. J.L.N. received funding from the NIH (CA225442). Grant funding to M.A.D. from the National Institutes of Health (CA245508, EY030180). The project was also supported by NIH grant CA21765 and American Lebanese Syrian Associated Charities.

## REFERENCES

1. Livesey, F.J. and C.L. Cepko, Vertebrate neural cell-fate determination: lessons from the retina. Nat Rev Neurosci, 2001. 2(2): p. 109–18.

2. Dyer, M.A. and C.L. Cepko, Regulating proliferation during retinal development. Nat Rev Neurosci, 2001. 2(5): p. 333–42.

3. Turner, D.L. and C.L. Cepko, A common progenitor for neurons and glia persists in rat retina late in development. Nature, 1987. 328(6126): p. 131-6.

4. Newman, E.A., Physiological properties and possible functions of Muller cells. Neurosci Res Suppl, 1986. 4(20): p. S209–20.

5. Newman, E. and A. Reichenbach, The Muller cell: a functional element of the retina. Trends Neurosci, 1996. 19(8): p. 307–12.

6. Bringmann, A. and A. Reichenbach, Heterogenous expression of Ca2+-dependent K+ currents by Muller glial cells. Neuroreport, 1997. 8(18): p. 3841–5.

7. Reichenbach, A., et al., The Muller (glial) cell in normal and diseased retina: a case for single-cell electrophysiology. Ophthalmic Res, 1997. 29(5): p. 326–40.

8. Bringmann, A., et al., Müller cells in the healthy and diseased retina. Prog Retin Eye Res, 2006. 25(4): p. 397–424.

9. Mata, N.L., et al., Isomerization and oxidation of vitamin a in cone-dominant retinas: a novel pathway for visual-pigment regeneration in daylight. Neuron, 2002. 36(1): p. 69–80.

10. Vecino, E., et al., Glia-neuron interactions in the mammalian retina. Prog Retin Eye Res, 2016. 51: p. 1–40.

11. Biswas, S., A. Cottarelli, and D. Agalliu, Neuronal and glial regulation of CNS angiogenesis and barriergenesis. Development, 2020. 147(9).

12. Goldman, D., Muller glial cell reprogramming and retina regeneration. Nat Rev Neurosci, 2014. 15(7): p. 431–42.

13. Lahne, M., et al., Reprogramming Müller Glia to Regenerate Retinal Neurons. Annu Rev Vis Sci, 2020. 6: p. 171–193.

14. Dyer, M.A. and C.L. Cepko, Control of Muller glial cell proliferation and activation following retinal injury. Nat Neurosci, 2000. 3(9): p. 873–80.

15. Fischer, A.J. and T.A. Reh, Muller glia are a potential source of neural regeneration in the postnatal chicken retina. Nat Neurosci, 2001. 4(3): p. 247–52.

16. Hoang, T., et al., Gene regulatory networks controlling vertebrate retinal regeneration. Science, 2020. 370(6519).

17. Aldiri, I., et al., The Dynamic Epigenetic Landscape of the Retina During Development, Reprogramming, and Tumorigenesis. Neuron, 2017. 94(3): p. 550–568 e10.

18. Norrie, J.L., et al., Nucleome Dynamics during Retinal Development. Neuron, 2019. 104(3): p. 512–528.e11.

19. Chen, B.C., et al., Lattice light-sheet microscopy: imaging molecules to embryos at high spatiotemporal resolution. Science, 2014. 346(6208): p. 1257998.

20. Dhingra, A., et al., Probing neurochemical structure and function of retinal ON bipolar cells with a transgenic mouse. The Journal of comparative neurology, 2008. 510(5): p. 484–96.

21. Huckfeldt, R.M., et al., Transient neurites of retinal horizontal cells exhibit columnar tiling via homotypic interactions. Nat Neurosci, 2009. 12(1): p. 35–43.

22. Siegert, S., et al., Genetic address book for retinal cell types. Nat Neurosci, 2009. 12(9): p. 1197–204.

23. Siegert, S., et al., Transcriptional code and disease map for adult retinal cell types. Nat Neurosci, 2012. 15(3): p. 487–95, S1-2.

24. Vazquez-Chona, F.R., A.M. Clark, and E.M. Levine, Rlbp1 promoter drives robust Muller glial GFP expression in transgenic mice. Investigative ophthalmology & visual science, 2009. 50(8): p. 3996–4003.

25. Masland, R.H., The neuronal organization of the retina. Neuron, 2012. 76(2): p. 266–80.

26. Chen, E.Y., et al., Enrichr: interactive and collaborative HTML5 gene list enrichment analysis tool. BMC Bioinformatics, 2013. 14: p. 128.

27. Dyer, M.A. and C.L. Cepko, p57(Kip2) regulates progenitor cell proliferation and amacrine interneuron development in the mouse retina. Development, 2000. 127(16): p. 3593–605.

28. Dyer, M.A. and C.L. Cepko, p27Kip1 and p57Kip2 regulate proliferation in distinct retinal progenitor cell populations. J. of Neurosci, 2001. 21: p. 4259–71.

29. Zhu, C.C., et al., Six3-mediated auto repression and eye development requires its interaction with members of the Groucho-related family of co-repressors. Development, 2002. 129(12): p. 2835–49.

30. Dyer, M.A., et al., Prox1 function controls progenitor cell proliferation and horizontal cell genesis in the mammalian retina. Nat Genet, 2003. 34(1): p. 53–8.

31. Donovan, S.L. and M.A. Dyer, Preparation and Square Wave Electroporation of Retinal Explant Cultures. Nature Protocols, 2006. 1(6): p. 2710–18.

32. McEvoy, J., et al., Coexpression of normally incompatible developmental pathways in retinoblastoma genesis. Cancer Cell, 2011. 20(2): p. 260–75.

33. Winkler, B.S., Relative inhibitory effects of ATP depletion, ouabain and calcium on retinal photoreceptors. Exp Eye Res, 1983. 36(4): p. 581–94.

34. Cutler, R.W., J.N. Young, and D.K. Urion, Factors influencing the vitreous potassium concentration in the rat. Invest Ophthalmol Vis Sci, 1983. 24(5): p. 631–6.

35. Mao, B.Q., P.R. MacLeish, and J.D. Victor, Role of hyperpolarization-activated currents for the intrinsic dynamics of isolated retinal neurons. Biophys J, 2003. 84(4): p. 2756–67.

36. Homma, K., et al., Developing rods transplanted into the degenerating retina of Crx-knockout mice exhibit neural activity similar to native photoreceptors. Stem Cells, 2013. 31(6): p. 1149–59.

37. Su, P.J., et al., Retinal synaptic regeneration via microfluidic guiding channels. Sci Rep, 2015. 5: p. 13591.

38. Rego, A.C., et al., Glutamate regulates the viability of retinal cells in culture. Vision Res, 2001. 41(7): p. 841–51.

39. Boccuni, I. and R. Fairless, Retinal Glutamate Neurotransmission: From Physiology to Pathophysiological Mechanisms of Retinal Ganglion Cell Degeneration. Life (Basel), 2022. 12(5).

40. Hoon, M., et al., Functional architecture of the retina: development and disease. Prog Retin Eye Res, 2014. 42: p. 44–84.

41. Hirasawa, H. and A. Kaneko, pH changes in the invaginating synaptic cleft mediate feedback from horizontal cells to cone photoreceptors by modulating Ca2+ channels. J Gen Physiol, 2003. 122(6): p. 657–71.

42. Beckwith-Cohen, B., et al., Localizing Proton-Mediated Inhibitory Feedback at the Retinal Horizontal Cell-Cone Synapse with Genetically-Encoded pH Probes. J Neurosci, 2019. 39(4): p. 651–662.

43. Zhu, D., et al., Regulation of vascular endothelial growth factor and pigment epithelium-derived factor in rat retinal explants under retinal acidification. Eye (Lond), 2009. 23(11): p. 2105–11.

44. Ham, Y., et al., Gitelman syndrome and ectopic calcification in the retina and joints. Clin Kidney J, 2021. 14(9): p. 2023–2028.

45. Marchini, G., et al., Choroidal calcification in Bartter syndrome. Am J Ophthalmol, 1998. 126(5): p. 727–9.

46. Schnichels, S., et al., Cyclosporine A Protects Retinal Explants against Hypoxia. Int J Mol Sci, 2021. 22(19).

47. Hirata, M., T.R. Shearer, and M. Azuma, Hypoxia Activates Calpains in the Nerve Fiber Layer of Monkey Retinal Explants. Invest Ophthalmol Vis Sci, 2015. 56(10): p. 6049–57.

48. Mei, S., et al., Mechanisms underlying somatostatin receptor 2 down-regulation of vascular endothelial growth factor expression in response to hypoxia in mouse retinal explants. J Pathol, 2012. 226(3): p. 519–33.

49. Natoli, R., et al., Expression and role of the early-response gene Oxr1 in the hyperoxia-challenged mouse retina. Invest Ophthalmol Vis Sci, 2008. 49(10): p. 4561–7.

50. Zhu, Y., et al., Microarray analysis of hyperoxia stressed mouse retina: differential gene expression in the inferior and superior region. Adv Exp Med Biol, 2010. 664: p. 217–22.

51. Yu, D.Y., S.J. Cringle, and E.N. Su, Intraretinal oxygen distribution in the monkey retina and the response to systemic hyperoxia. Invest Ophthalmol Vis Sci, 2005. 46(12): p. 4728–33.

52. Chen, G.H. and C.C. Chiao, Mild stress culture conditions promote neurite outgrowth of retinal explants from postnatal mice. Brain Res, 2020. 1747: p. 147050.

53. Caprioli, J., S. Kitano, and J.E. Morgan, Hyperthermia and hypoxia increase tolerance of retinal ganglion cells to anoxia and excitotoxicity. Invest Ophthalmol Vis Sci, 1996. 37(12): p. 2376–81.

54. Tytell, M., M.F. Barbe, and I.R. Brown, Induction of heat shock (stress) protein 70 and its mRNA in the normal and light-damaged rat retina after whole body hyperthermia. J Neurosci Res, 1994. 38(1): p. 19–31.

55. Maliha, A.M., et al., Diminished apoptosis in hypoxic porcine retina explant cultures through hypothermia. Sci Rep, 2019. 9(1): p. 4898.

56. Schultheiss, M., et al., Hypothermia Protects and Prolongs the Tolerance Time of Retinal Ganglion Cells against Ischemia. PLoS One, 2016. 11(2): p. e0148616.

57. Rey-Funes, M., et al., Hypothermia prevents gliosis and angiogenesis development in an experimental model of ischemic proliferative retinopathy. Invest Ophthalmol Vis Sci, 2013. 54(4): p. 2836–46.

58. Liberski, S., B.J. Kaluzny, and J. Kocięcki, Methanol-induced optic neuropathy: a still-present problem. Arch Toxicol, 2022. 96(2): p. 431–451.

59. Liu, D.M., et al., The Intoxication Effects of Methanol and Formic Acid on Rat Retina Function. J Ophthalmol, 2016. 2016: p. 4087096.

60. Eells, J.T., et al., Development and characterization of a rodent model of methanol-induced retinal and optic nerve toxicity. Neurotoxicology, 2000. 21(3): p. 321–30.

61. Liesivuori, J. and H. Savolainen, Methanol and formic acid toxicity: biochemical mechanisms. Pharmacol Toxicol, 1991. 69(3): p. 157–63.

62. Broderick, C., et al., IFN-gamma and LPS-mediated IL-10-dependent suppression of retinal microglial activation. Invest Ophthalmol Vis Sci, 2000. 41(9): p. 2613–22.

63. Ghosh, F., et al., Retinal neuroinflammatory induced neuronal degeneration – Role of toll-like receptor-4 and relationship with gliosis. Exp Eye Res, 2018. 169: p. 99–110.

64. Carter, D.A. and A.D. Dick, Lipopolysaccharide/interferon-gamma and not transforming growth factor beta inhibits retinal microglial migration from retinal explant. Br J Ophthalmol, 2003. 87(4): p. 481–7.

65. Maldonado, R.F., I. Sá-Correia, and M.A. Valvano, Lipopolysaccharide modification in Gram-negative bacteria during chronic infection. FEMS Microbiol Rev, 2016. 40(4): p. 480–93.

66. Geiger, K., et al., Transgenic mice expressing IFN-gamma in the retina develop inflammation of the eye and photoreceptor loss. Invest Ophthalmol Vis Sci, 1994. 35(6): p. 2667–81.

67. Husain, S., et al., Interferon-gamma (IFN-γ)-mediated retinal ganglion cell death in human tyrosinase T cell receptor transgenic mouse. PLoS One, 2014. 9(2): p. e89392.

68. London, A., I. Benhar, and M. Schwartz, The retina as a window to the brain-from eye research to CNS disorders. Nat Rev Neurol, 2013. 9(1): p. 44–53.

69. Singh, S., S. Singh, and A. Kumar, Systemic Candida albicans Infection in Mice Causes Endogenous Endophthalmitis via Breaching the Outer Blood-Retinal Barrier. Microbiol Spectr, 2022. 10(4): p. e0165822.

70. Fan, K.C., et al., In Vitro Susceptibilities of Vitreous Candida Endophthalmitis Isolates to Newer and Traditional Antifungal Agents. Ophthalmic Surg Lasers Imaging Retina, 2019. 50(5): p. S13–s17.

71. Klotz, S.A., et al., Fungal and parasitic infections of the eye. Clin Microbiol Rev, 2000. 13(4): p. 662–85.

72. Shah, C.P., et al., Ocular candidiasis: a review. Br J Ophthalmol, 2008. 92(4): p. 466–8.

73. Ksiaa, I., et al., Update on Bartonella neuroretinitis. J Curr Ophthalmol, 2019. 31(3): p. 254–61.

74. Lundquist, O. and S. Osterlin, Glucose concentration in the vitreous of nondiabetic and diabetic human eyes. Graefes Arch Clin Exp Ophthalmol, 1994. 232(2): p. 71–4.

75. Fu, J., et al., Essential Functions of the Transcription Factor Npas4 in Neural Circuit Development, Plasticity, and Diseases. Front Neurosci, 2020. 14: p. 603373.

76. Li, Y., et al., A critical evaluation of the activity-regulated cytoskeleton-associated protein (Arc/Arg3.1)’s putative role in regulating dendritic plasticity, cognitive processes, and mood in animal models of depression. Front Neurosci, 2015. 9: p. 279.

77. Yakout, D.W., N. Shree, and A.M. Mabb, Effect of pharmacological manipulations on Arc function. Curr Res Pharmacol Drug Discov, 2021. 2: p. 100013.

78. Jia, L., et al., The roles of TNFAIP2 in cancers and infectious diseases. J Cell Mol Med, 2018. 22(11): p. 5188–5195.

79. Lipson, K.E., et al., CTGF is a central mediator of tissue remodeling and fibrosis and its inhibition can reverse the process of fibrosis. Fibrogenesis Tissue Repair, 2012. 5(Suppl 1): p. S24.

80. Hamon, A., et al., Retinal Degeneration Triggers the Activation of YAP/TEAD in Reactive Müller Cells. Invest Ophthalmol Vis Sci, 2017. 58(4): p. 1941–1953.

81. Chen, Y., et al., Regulations of Retinal Inflammation: Focusing on Müller Glia. Front Cell Dev Biol, 2022. 10: p. 898652.

82. Rosenbaum, J.T., et al., Endotoxin-induced uveitis in rats as a model for human disease. Nature, 1980. 286(5773): p. 611-3.

83. Miserocchi, E., et al., Review on the worldwide epidemiology of uveitis. Eur J Ophthalmol, 2013. 23(5): p. 705–17.

84. Miyamoto, K., et al., In vivo quantification of leukocyte behavior in the retina during endotoxin-induced uveitis. Invest Ophthalmol Vis Sci, 1996. 37(13): p. 2708–15.

85. Koga, T., et al., Coinduction of nitric oxide synthase and arginine metabolic enzymes in endotoxin-induced uveitis rats. Exp Eye Res, 2002. 75(6): p. 659–67.

86. de Vos, A.F., V.N. Klaren, and A. Kijlstra, Expression of multiple cytokines and IL-1RA in the uvea and retina during endotoxin-induced uveitis in the rat. Invest Ophthalmol Vis Sci, 1994. 35(11): p. 3873–83.

87. Shen, D.F., et al., Cytokine gene expression in different strains of mice with endotoxin-induced uveitis (EIU). Ocul Immunol Inflamm, 2000. 8(4): p. 221–5.

88. Koizumi, K., et al., Contribution of TNF-alpha to leukocyte adhesion, vascular leakage, and apoptotic cell death in endotoxin-induced uveitis in vivo. Invest Ophthalmol Vis Sci, 2003. 44(5): p. 2184–91.

89. Cousins, S.W., et al., Endotoxin-induced uveitis in the rat: observations on altered vascular permeability, clinical findings, and histology. Exp Eye Res, 1984. 39(5): p. 665–76.

90. Okumura, A., et al., Endotoxin-induced uveitis (EIU) in the rat: a study of inflammatory and immunological mechanisms. Int Ophthalmol, 1990. 14(1): p. 31–6.

91. McMenamin, P.G. and J. Crewe, Endotoxin-induced uveitis. Kinetics and phenotype of the inflammatory cell infiltrate and the response of the resident tissue macrophages and dendritic cells in the iris and ciliary body. Invest Ophthalmol Vis Sci, 1995. 36(10): p. 1949–59.

92. Kurihara, T., et al., Neuroprotective effects of angiotensin II type 1 receptor (AT1R) blocker, telmisartan, via modulating AT1R and AT2R signaling in retinal inflammation. Invest Ophthalmol Vis Sci, 2006. 47(12): p. 5545–52.

93. Zhuang, X., et al., SHP-1 suppresses endotoxin-induced uveitis by inhibiting the TAK1/JNK pathway. J Cell Mol Med, 2021. 25(1): p. 147–160.

94. Du, L., et al., Ruxolitinib Alleviates Uveitis Caused by Salmonella typhimurium Endotoxin. Microorganisms, 2021. 9(7).

95. Huang, Q., et al., Evaluation of Cell Type Annotation R Packages on Single-cell RNA-seq Data. Genomics Proteomics Bioinformatics, 2021. 19(2): p. 267–281.

96. Mashimo, H., et al., Neutrophil chemotaxis and local expression of interleukin-10 in the tolerance of endotoxin-induced uveitis. Invest Ophthalmol Vis Sci, 2008. 49(12): p. 5450–7.

97. Hughes, C.E. and R.J.B. Nibbs, A guide to chemokines and their receptors. Febs j, 2018. 285(16): p. 2944–2971.

98. Zlotnik, A. and O. Yoshie, The chemokine superfamily revisited. Immunity, 2012. 36(5): p. 705–16.

99. Nourshargh, S., P.L. Hordijk, and M. Sixt, Breaching multiple barriers: leukocyte motility through venular walls and the interstitium. Nat Rev Mol Cell Biol, 2010. 11(5): p. 366–78.

100. Capucetti, A., F. Albano, and R. Bonecchi, Multiple Roles for Chemokines in Neutrophil Biology. Front Immunol, 2020. 11: p. 1259.

101. Goldman, D., Müller glial cell reprogramming and retina regeneration. Nat Rev Neurosci, 2014. 15(7): p. 431–42.

102. Nomiyama, H., N. Osada, and O. Yoshie, The evolution of mammalian chemokine genes. Cytokine Growth Factor Rev, 2010. 21(4): p. 253–62.

103. DeVries, M.E., et al., Defining the origins and evolution of the chemokine/chemokine receptor system. J Immunol, 2006. 176(1): p. 401–15.

104. Sommer, F., V. Torraca, and A.H. Meijer, Chemokine Receptors and Phagocyte Biology in Zebrafish. Front Immunol, 2020. 11: p. 325.

105. Jorstad, N.L., et al., Stimulation of functional neuronal regeneration from Muller glia in adult mice. Nature, 2017. 548(7665): p. 103-107.

