## supplemental figures and methods for "Latent Epigenetic Programs in Müller Glia Contribute to Stress, Injury, and Disease Response in the Retina"

**SUPPLEMENTAL METHODS**

**WGBS, ChIP-seq, and ATAC-seq**

DNA methylation (whole-genome bisulfite sequencing, WGBS), histone modifications and transcriptional regulation through enhancer interactions (ChIP-seq on histone marks, Brd4 and RNA polymerase II, Pol II) and chromatin accessibility (assay for transposase-accessible chromatin using sequencing, ATAC-seq) across retinal development was previously reported and analyzed (Aldiri et al. 2017)**.** Additional retina treated for 24 hours with NMDA or LPS were processed as previously reported (Aldiri et al. 2017)**.**

**Human and mouse retinal explant cultures**

All animal procedures and protocols were approved by the St. Jude Laboratory Animal Care and Use Committee under protocol number 393-100500. All studies conform to federal and local regulatory standards. Mice were housed on ventilated racks on a standard 12 h light–dark cycle.

**Human Retina were obtained from the Midsouth Eyebank.**

Retina were dissected from the eye cup in Retinal Explant Media (REM). 1.5mm punches were taken from each eye using a biopsy punch (Integra #33-31A-P/25). Punches were placed on a Nuclepore 1.0uM membrane (Whatman #110410) floating on 2mL of REM (45% Hams F-12, 45% DMEM, 10%FBS, 10mL/L HEPES, 5mg/mL Insulin, 10mL/L Penicillin-Streptomycin-Glutamine) in a 12-well dish. 50uL of REM was removed from the dish and placed on the retinal punches. The retinal punches were incubated for 24 hours (37C, 5% CO2) for culture control retina. The morning of harvest an additional 50uL of REM was added to the floating retinal punches to ensure they would not dry out. Punches were harvested after 24 hours.

Retinal punches were stressed by adding a stressor to the REM or by culturing in different conditions listed below:

| **Sample** | **Temp (°C)** | **Oxygen (%)** | **Treatment** |
| --- | --- | --- | --- |
| *B. henselae* | 37 | 20 | 100 CFU/mL |
| 33°C | 33 | 20 |  |
| 40°C | 40 | 20 |  |
| *C. albicans* | 37 | 20 | 100 CFU/mL |
| control culture | 37 | 20 |  |
| glucose | 37 | 20 | 20 mM glucose |
| glutamate | 37 | 20 | 1mM glutamate |
| IFN | 37 | 20 | 2ul IFNg (100U/mL) |
| KCl | 37 | 20 | 80 mM KCl |
| LPS | 37 | 20 | 0.2mg/mL LPS |
| MeOH | 37 | 20 | 8 mM formic acid |
| 2% O2 | 37 | 2.3 |  |
| 40% O2 | 37 | 40 |  |
| ouabain | 37 | 20 | 70 mM ouabain |
| pH 7.15 | 37 | 20 | HCl to pH 7.15 |
| pH 7.7 | 37 | 20 | NaOH to pH 7.7 |

**Immunoblots**

For cytokine array panel, retina were dissected and the corners were cut to lay flat the nucleopore membrane. Cultures were completed as described above with 1 retina per membrane and 100uL of media added to the top of these cultures. Retina were cultured for 24 hours. Two retina per condition were transferred to 1mL RIPA buffer and protein was extracted using a pestle to homogenize the tissue. 1mL of each sample was added to a pre-blocked membrane per protocol of the Mouse Cytokine Array Panel A (R&D Systems #ARY006). Array was completed according to the manufacture’s protocol.

**Dissociation of retina and retinal punches for single cell sequencing**

Papatin buffer was prepared by adding L-cysteine (1mM final concentration) and EDTA (0.5mM final concentration) to PBS -/-. Then 40U papain (Worthington CAT#LS003119) was added to 400uL of papain buffer and incubate at 37°C for ~10 min. Retina or retinal punches were added to 400uL of papain and incubated at 37°C. The tubes were flicked every 2-5 minutes until the tissue was dissociated. 40 μl of Dnase was added and the tube was incubated for an additional 5 min in a 37 °C water bath. The suspension was transferred into a 50-ml conical tube through a 40-μm mesh cell strainer and washed with PBS^−/−^ to bring the total volume to 3 ml. A 5 ml BSA cushion medium (4% BSA in Retinal Explant media (or any media with 10% serum)) was prepared in a 15-ml conical tube and the filtered cell suspension was slowly layered on top of the cushion. The sample was centrifuged at 500*g* for 10 min at 4 °C and the cell pellet was resuspended in REM. Cells were counted and viability was confirmed on a hemocytometer.

**ScRNAseq**

Approximately 10,000 cells from each sample were taken and loaded onto the 10× chromium controller for single-cell RNA-sequencing analysis which was completed according to the 10× genomics protocol. Barcoded sequences were sequenced according to 10× Genomics protocol on an Illumina HiSeq 2500 or 4000.

**ScATACseq**

Nuclei were extracted from retina following the Nuclei Isolation from Embryonic Mouse Brain for Single Cell Multiome ATAC + Gene Expression Sequence demonstrated protocol from 10x genomics. Briefly, retinal cells dissociated in papain were resuspended in 0.1X Lysis Buffer (10mM Tris-Hcl 7.4, 10mM NaCl, 3mM MgCl_2_, 1% BSA, 1mM DTT, 1U/uL Rnase Inhibitor, 0.01% Tween-20, 0.01% NP40, 0.001% Digitonin) for 2 minutes on ice. 1ML Wash buffer was added (10mM Tris-Hcl 7.4, 10mM NaCl, 3mM MgCl_2_, 1% BSA, 1mM DTT, 1U/uL Rnase Inhibitor, 0.01% Tween-20) and the nuclei were spun down at 500g for 5 min. The supernatent was removed and the nuclei was resuspended in 1x Nuceli Buffer (10x Genomics 2000153) with 1U/uL RNase inhibitor. ScATAC-seq was completed with using the 10x genomics Chromium Next GEM Single Cell ATAC Reagent Kits (PN-1000390). Barcoded sequences were sequenced according to 10× Genomics protocol on an Illumina HiSeq 2500 or 4000.

**Bulk RNAseq**

**RNA extractions for retinal punches**

Retinal punches were dissolved in 800uL Trizol Reagent (Invitrogen #15596026). 200uL chloroform was added to each sample, inverted, incubated at room temperature for 3 minutes. Samples were then spun down at 12,000 3 g at 4 C for 15 min. The aqueous later was transferred to a new tube and 1uL glycogen and 500 uL isopropanol was added to each sample and mixed by vortex. Samples were incubated at room temperature for 10 minutes and spun down at 12,000 3 g at 4 C for 15 min. The isopropanol was removed and the pellet was washed twice with ice-cold 80% EtOH. The washed pellet was resuspened in water.

**RNA-sequencing and analysis**

RNA-sequencing libraries were prepared with the TruSeq Stranded Total RNA Library Prep Kit and paired-end sequencing was performed on HiSeq2000 or 2500 sequencers (Illumina). Reads were aligned using STAR Aligner (v2.7) (mm10 for mouse or hg38 for human) and gene quantification was determined using RSEM (v1.3.1). The top 5000 genes were used for a Principal Component Analysis (PCA).

**qPCR**

Retina were dissolved in 800uL Trizol Reagent (Invitrogen #15596026). 400 uL of the Trizol was used for RNA extraction using the Zymo Direct-Zol Miniprep Kit (Zymo #R2052). cDNA was created with 150ng RNA using a High-Capacity RNA-to-cDNA™ Kit (Applied Biosystems 4387406). qPCR was run using SYBR Select Master Mix(Applied Biosystems 4472908) according to the manufacture’s protocol on a QuantStudio3 (Applied Biosystems).

Cxcl2_F AGTGAACTGCGCTGTCAATG

Cxcl2_R TTCAGGGTCAAGGCAAACTT

Ccl5_F CTGCTGCTTTGCCTACCTCT

Ccl5_R ACACACTTGGCGGTTCCTT

Il6_F CCGGAGAGGAGACTTCACAG

Il6_R TTCTGCAAGTGCATCATCGT

Gapdh_F CTCCACTCACGGCAAATTCA

Gapdh_R CGCTCCTGGAAGATGGTGAT

**In-vivo LPS**

LPS was reconstituted at 1mg/mL in PBS -/-. Prior to injection LPS was further diluted in PBS-/- to a concentration of 25ng/uL. Mice were anesthetized with isoflurane and a small puncture was created at the edge of the sclera with a needle after which a hamilton syringe was used to inject 1uL into the posterior chamber of the eye.After 24 hours eyes were dissected in 100uL PBS -/-. The retina was removed and added to 800uL Trizol.

The 100uL of vitreous/PBS -/- used for dissection is transferred to a flow cytometry tube.

**Flow Cytometry Analysis of Vitreous**

**CD45 and AccuCount Beads**

Blocking antibody (1:100 eBiosciences Anti-mouse CD16/32 16-0161-86) was added to each sample, vortexed and incubated at room temperature at 5 min. 50uL of antibody and bead master mix were added to each sample (3uL antibody (CD45, BioLegend 30-F11), 20uL beads (AccuCount 7.9um, Spherotech ACRP-70-5), and 27uL PBS-/-). Samples were incubated for 12 min in the dark, spun down for 5 min at 500g. Supernatant was poured off and pellet was resuspended in 150uL PBS. Cells were analyzed on a BioRad S3e Cell Sorter.

**Full Immune Panel**

Splenocytes and bone marrow from 1 wildtype murine spleen was used as a control. Red blood cells were lysed per manufacture’s protocol. Two million cells were aliquoted per control tubes and spun down. Cells were washed twice with 500uL PBS and resuspended in 200uL Live/Dead Blue Mix and incubated on ice for 30 minutes. Add 100uL of staining media (1X HBSS; 2% (v/v) calf serum; 10mM NaN3, 10 mM HEPES [pH 7.2]), vortex, spin down, and decant. Resuspend each tube in 100uL of block, incubated 30 minute on ice, add 100uL of staining media, vortex, spin down, and decant. Resuspend cell pellets in 75uL of primary stain mix, incubate on 30 minutes on ice, add 500uL staining media, vortex, spin down and decant. Resuspend final pellets in 200uL staining media.

| **Antibodies** | **Dilution** |
| --- | --- |
| CD45 (30-F11)-BUV395 | 200 |
| CD19 (6D5)-BUV661 | 200 |
| CD3 (500A2)-BV480 | 50 |
| Ly6G-BV605 | 100 |
| MHCII BV786 | 200 |
| CD68-FITC | 100 |
| CD11b-PerCP-Cy5.5 | 100 |
| CD11c-PE | 100 |
| CD49b-PE-Cy5.5 | 100 |
| Siglec-F-APC | 100 |
| Ly6C-Alexa700 | 100 |
| NK1.1-APC-Cy7 | 100 |

**Knockout Mouse**

Cxcr2 knockout mice (Strain #:006848) were purchased from Jackson Laboratory.

Mice were genotyped using the following primers:

*Cxcr2 WT-F:* 5’-AGAATTGGGGTCAGGGTACA-3’

*Cxcr2 WT-R:* 5’-CCCTTAGGCTGCAAATGAAC-3’

*Cxcr2 mut-F:* 5’-CTTGGGTGGAGAGGCTATTC-3’

*Cxcr2 mut-R:* 5’-AGGTGAGATGACAGGAGATC-3’

PCR PROGRAM for Cxcr2

Denature: 94°C 2’

Cycle (10): 94°C 20’’, 65-->60°C 15’’, 68°C 10’’

Cycle (28): 94°C 15’’, 60°C 15’’, 72°C 10’’

Final extension: 72°C 2’

Hold: 4°C

EXPECTED PRODUCT SIZE: KO 280 bp; WT 214 bp

**Lattice Light Sheet Microscope (LLSM) Imaging:**

LLSM cover slip preparation: 5mm glass cover slips were cleaned by treatment with 1M NaOH, followed by washing in water and sterilization in EtOH. Cover slips were coated in Poly-L-Lysin (Sigma, diluted 1:10 in PBS) for 30 min at 37 deg, and washed 2x with PBS before plating cells.

Retina were dissociated as described above and ~120 ul plated on a chamber slide containing 5mm glass coverslips for imaging.

LLSM cover slip preparation: 5mm glass cover slips were cleaned by treatment with 1M NaOH, followed by washing in water and sterilization in EtOH. Cover slips were coated in Poly-L-Lysin (Sigma, diluted 1:10 in PBS) for 30 min at 37 deg, and washed 2x with PBS before plating cells.

The Lattice Light Sheet Microscope (3i-Intellegent Imaging Innovations) is equipped with a 25x dipping lens, with a 1.1 NA.

Nuclear Probe Validation: Imaging was performed with a 12um sheet (delta NA .5-.411) and a square pattern. Cells were acquired in dither mode using sample scan at a step size of .3um, resulting in voxels 104nm in x and y and 163nm in z. SNAP and HALO tag constructs were labelled with JF dyes from the Lavis Lab at the Howard Hughes Medical institute^1^.

Nuclear Classifier: Imaging was performed with a 7.5um sheet (delta NA .5-.38) and a hexagonal pattern. Cells were acquired in the same mode as previously stated. Cells from each transgenic mouse line were stained during plating with 1uM Sir-640 dye (Spirochrome cat#sc007) for DNA visualization.

**Cell-type classification model**

We used the supervised auto-context machine-learning method in ilastik^2^, an open-source image analysis software package, to segment the 3D images of nuclei based on intensities into three classes, high, mid and low that correspond to different DNA densities. We used Fiji to measure 3D volume, surface area and morphometric properties of each region in the nucleus^3^. We computed Haralick texture features using mahotas library^4^ for each nuclei. We combined the two feature sets into a feature matrix each row representing a cell and columns representing the features. We randomly sampled the rows per cell type and used 66% of the samples as the training set and used the rest for validation. We used the training data to train a random forest machine-learning algorithm to predict the cell type in the validation dataset. We created 2000 randomly assigned training and validation dataset splits. For each pair of datasets, we trained a random-forest model in H2O machine learning R package (version 3.20.0.8) and recorded the probability predictions of the model for the validation datasets for each class. We calculate the mean of the probabilities per class for each image and called the cell type based on the highest mean probability among the five classes. We calculate the maximum of the mean probabilities and used it as a measure of confidence for the model when the prediction is wrong. This allowed us to define prediction a probability threshold per class above which model predictions are precise.

**Bone Marrow Transplant**

Donor cell isolation:

Bone marrow was harvested from donor mouse femurs by flushing with PBS and spun down at 500g x 5 min. after removing the supernatant, cells were treated by adding 2ml of Puregene RBC lysis solution (Qiagen cat #158106) and incubated at RT for 10 min. After incubation, 8 ml of PBS/10%FBS was added to the tube, and cells were spun again at 500gx5min. Cells were resuspended in RPMI at 50e6 cells/ml and kept on ice till injection.

Recipient preparation and transplant:

At 4 weeks of age, mice were lethally irradiated to 9 Gy in an Xstrahl CIX3-irradiation cabinet using a Thoraeus filter. Within 4 hours of treatment, 5e6 cells were injected by tail vein. Mice were given injectable Baytril prophylactically by subcutaneous injection (5mg/kg) for three days before irradiation, and given Baytril treated water for 7 days after and were monitored for weight loss.

1	Grimm, J. B., Brown, T. A., English, B. P., Lionnet, T. & Lavis, L. D. Synthesis of Janelia Fluor HaloTag and SNAP-Tag Ligands and Their Use in Cellular Imaging Experiments. *Methods Mol Biol* **1663**, 179-188 (2017). <https://doi.org:10.1007/978-1-4939-7265-4_15>

2	Berg, S. *et al.* ilastik: interactive machine learning for (bio)image analysis. *Nat Methods* **16**, 1226-1232 (2019). <https://doi.org:10.1038/s41592-019-0582-9>

3	Schindelin, J. *et al.* Fiji: an open-source platform for biological-image analysis. *Nat Methods* **9**, 676-682 (2012). <https://doi.org:10.1038/nmeth.2019>

4	Coelho, Luis Pedro. "Mahotas: Open source software for scriptable computer vision." *arXiv preprint arXiv:1211.4907* (2012).

**SUPPLEMENTAL FIGURES**

**Figure S1. Retinal cell types can be identified by their nuclear morphology. A)** Violin plot of nuclear volume for euchromatin, facultative heterochromatin and constitutive heterochromatin across 5 classes of retinal cell types. **B)** Bar plot for confidence score for the machine learning based algorithm to identify retinal cell types based on nuclear features alone.

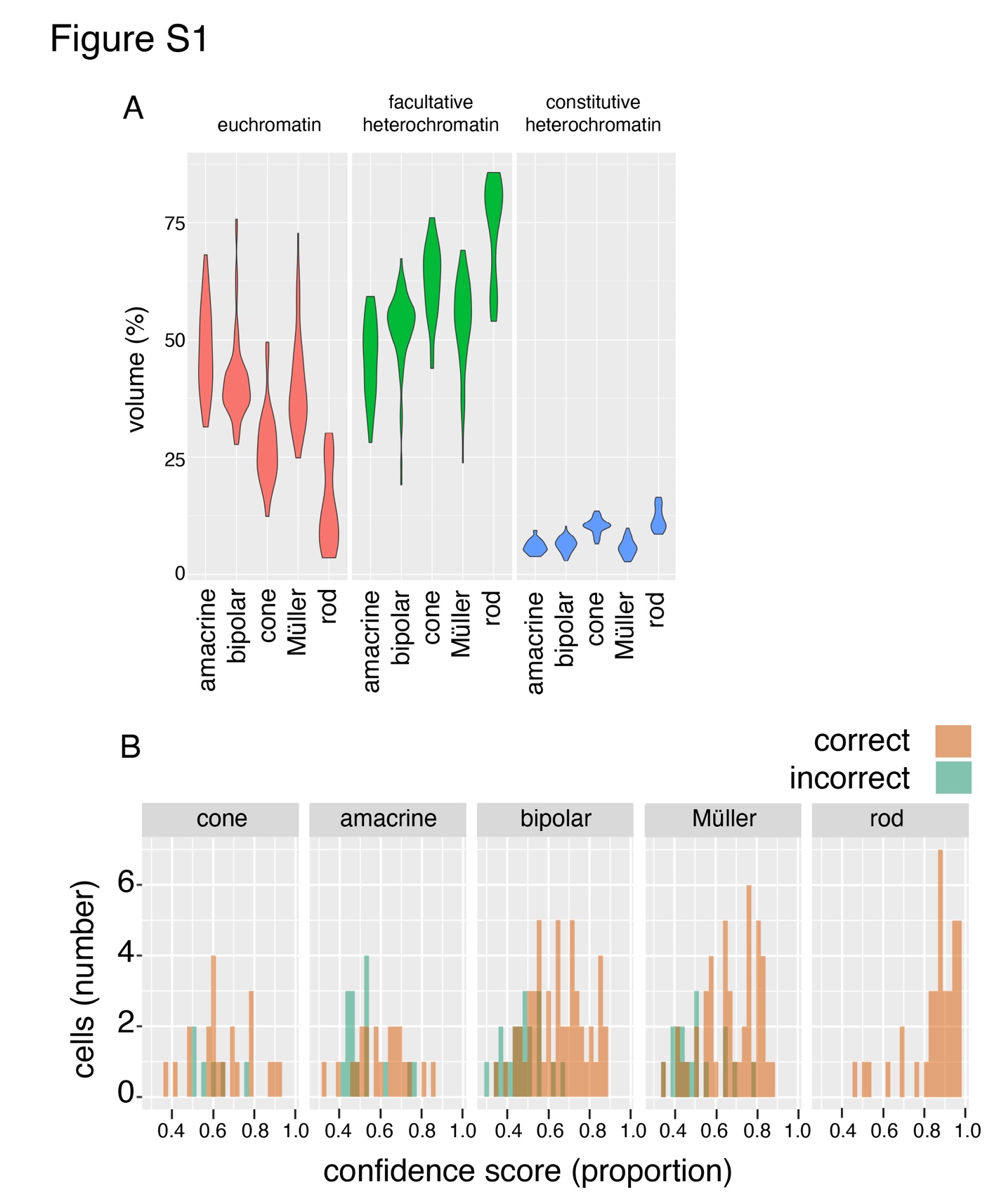

**Figure S2. Cell-type specific promoter chromatin accessibility is a feature of cell-type specific gene expression. A)** DNA methylation (WGBS), bulk ATAC-seq and ChIP-seq tracks for the *Aipl1* gene at E14.5. The gene is not expressed (barplot at bottom of figure) and the promoter is hypermethylated. **B)** The same tracks in the adult retina show reduced DNA methylation, increased promoter accessibility in bulk ATAC-seq and ChIP-seq profiles consistent with gene expression at this stage. **C)** Tracks for scATAC-seq showing specific chromatin accessibility in rods and cones (arrows) that express the gene. **D)** DNA methylation, bulk ATAC-seq and ChIP-seq for the *Sfrp1* gene at E14.5 when it is expressed (see barplot at bottom of figure). **E)** In the adult retina, the gene is silenced by increased H3K27me3 (arrow). **F)** The progenitor cell expression is correlated with elevated promoter accessibility in retinal progenitor cells and neurogenic (newly postmitotic) cells at E14.5 (arrows). Abbreviations: WGBS, whole genome bisulfite sequencing; kb, kilobase; FPKM, fragments per kilobase per million reads.

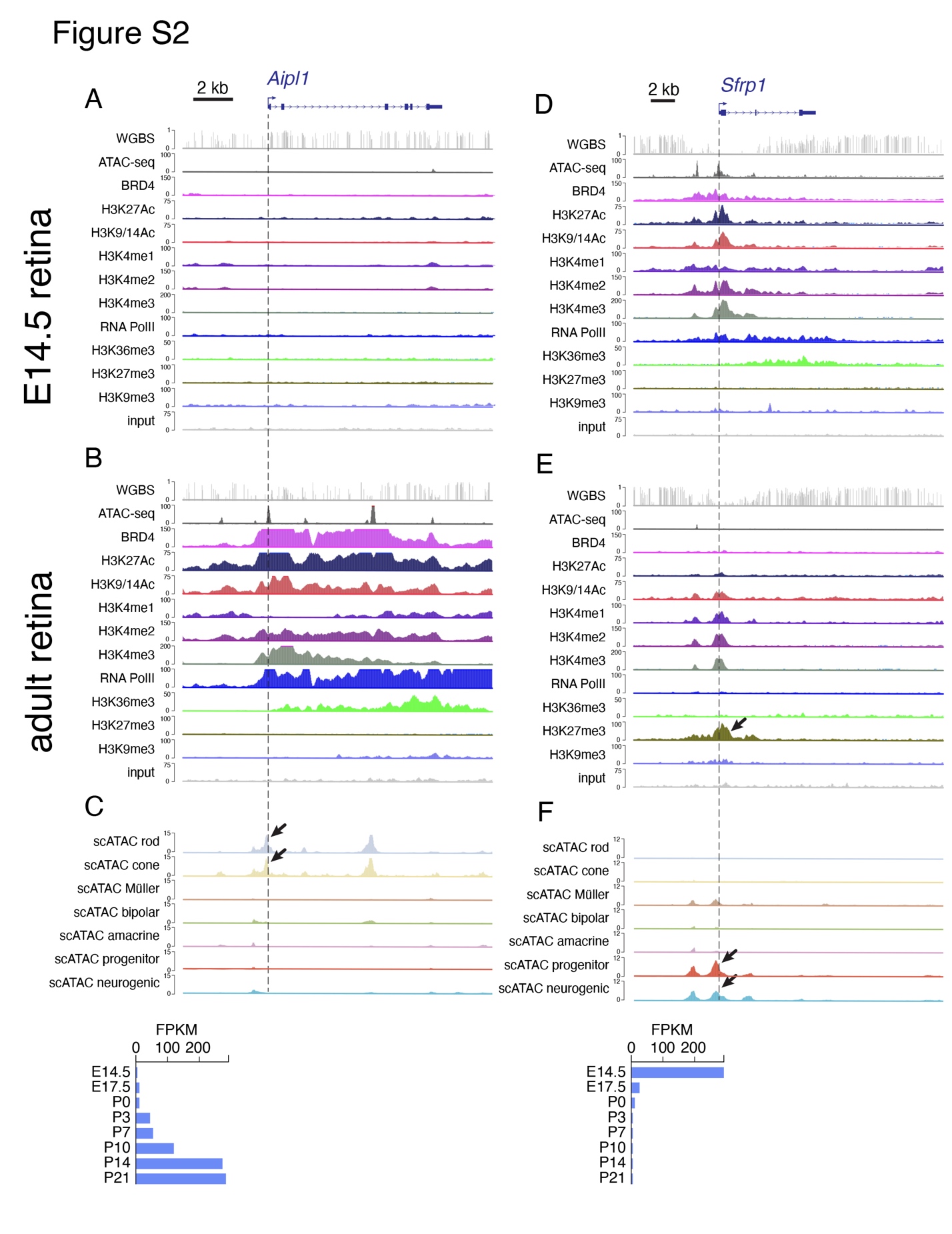

**Figure S3. Correlation between scRNA-seq and scATAC-seq. A)** DNA methylation (WGBS), bulk ATAC-seq and ChIP-seq tracks for the *Abca4* gene in the adult retina. The gene is expressed in rods and cones and increased during differentiation (barplot at right). **B)** Tracks for scATAC-seq showing specific chromatin accessibility in rods and cones (arrows) that express the gene. **C)** Barplots for scRNA-seq and scATAC-seq showing the expression (FPKM) in each cell population, percent of cells expressing the gene in each cell population, chromatin accessibility (RPKM) and percent of cells with accessible chromatin in each population. Abbreviations: WGBS, whole genome bisulfite sequencing; kb, kilobase; FPKM, fragments per kilobase per million reads; RPKM, reads per kilobase per million reads.

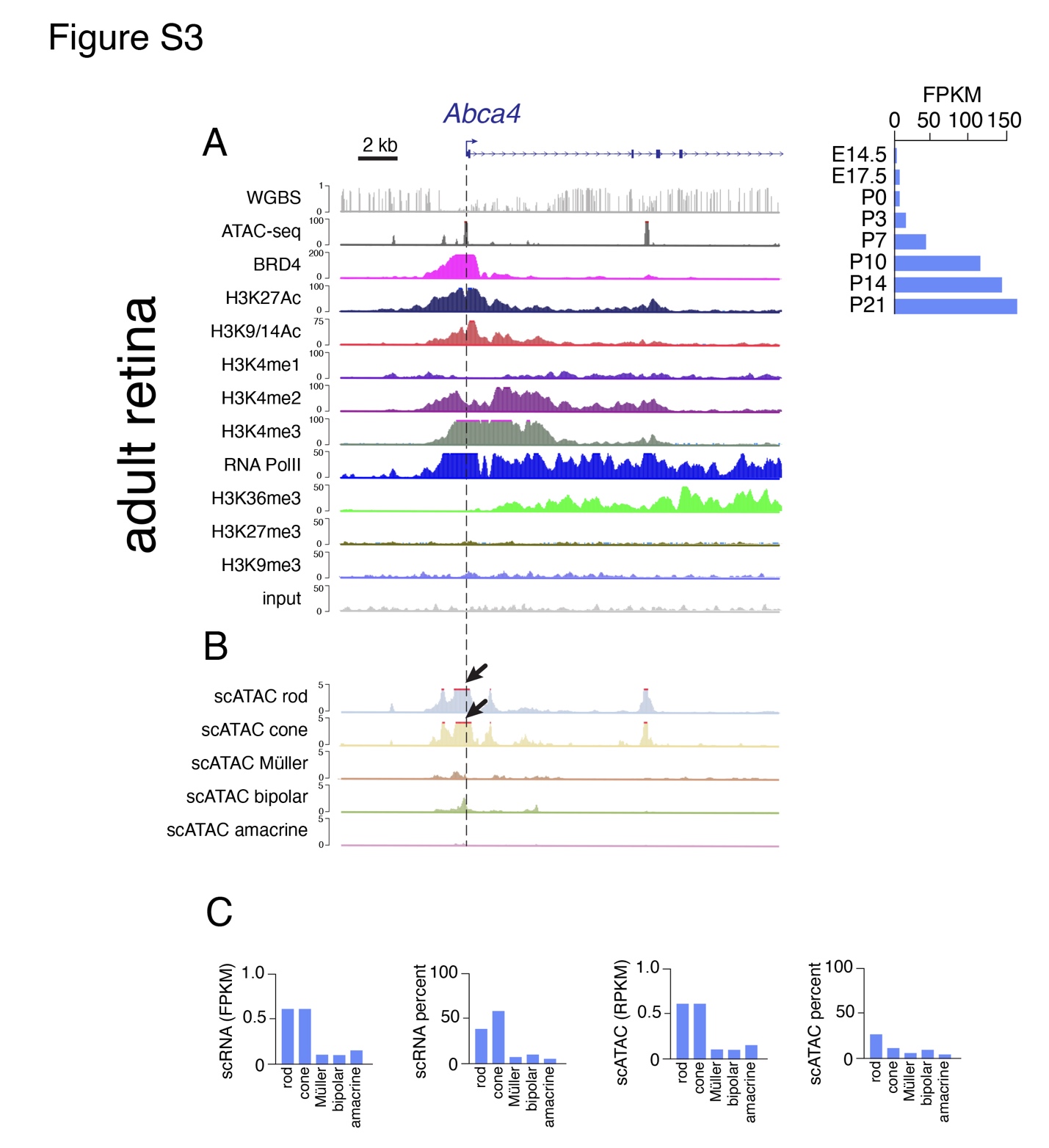

**Figure S4. Pliancy genes have a unique chromatin landscape. A)** DNA methylation, bulk ATAC-seq, ChIP-seq and scATAC-seq for *Cxcl12* in the E14.5 mouse retina. The expression across development in bulk RNA-seq is shown in the barplot at the right. **B)** Similar tracks are shown for the adult retina under normal conditions. The arrow highlight the lack of RNA-polII ChIP-seq signal at the promoter consistent with a lack of expression. However, there is a peak at the promoter in scATAC-seq in Müller glia (arrow). **C)** 24 hours after exposure to NMDA or LPS, there is a dramatic increase in ChIP-seq for RNA-polII ChIP-seq consistent with the increase expression in bulk RNA-seq. **D,E)** Single cell RNA-seq of human (D) or mouse (E) retina following LPS exposure shows increase expression is primarily in the Müller glia. Abbreviations: WGBS, whole genome bisulfite sequencing; kb, kilobase; UMAP, uniform manifold approximation and projection.

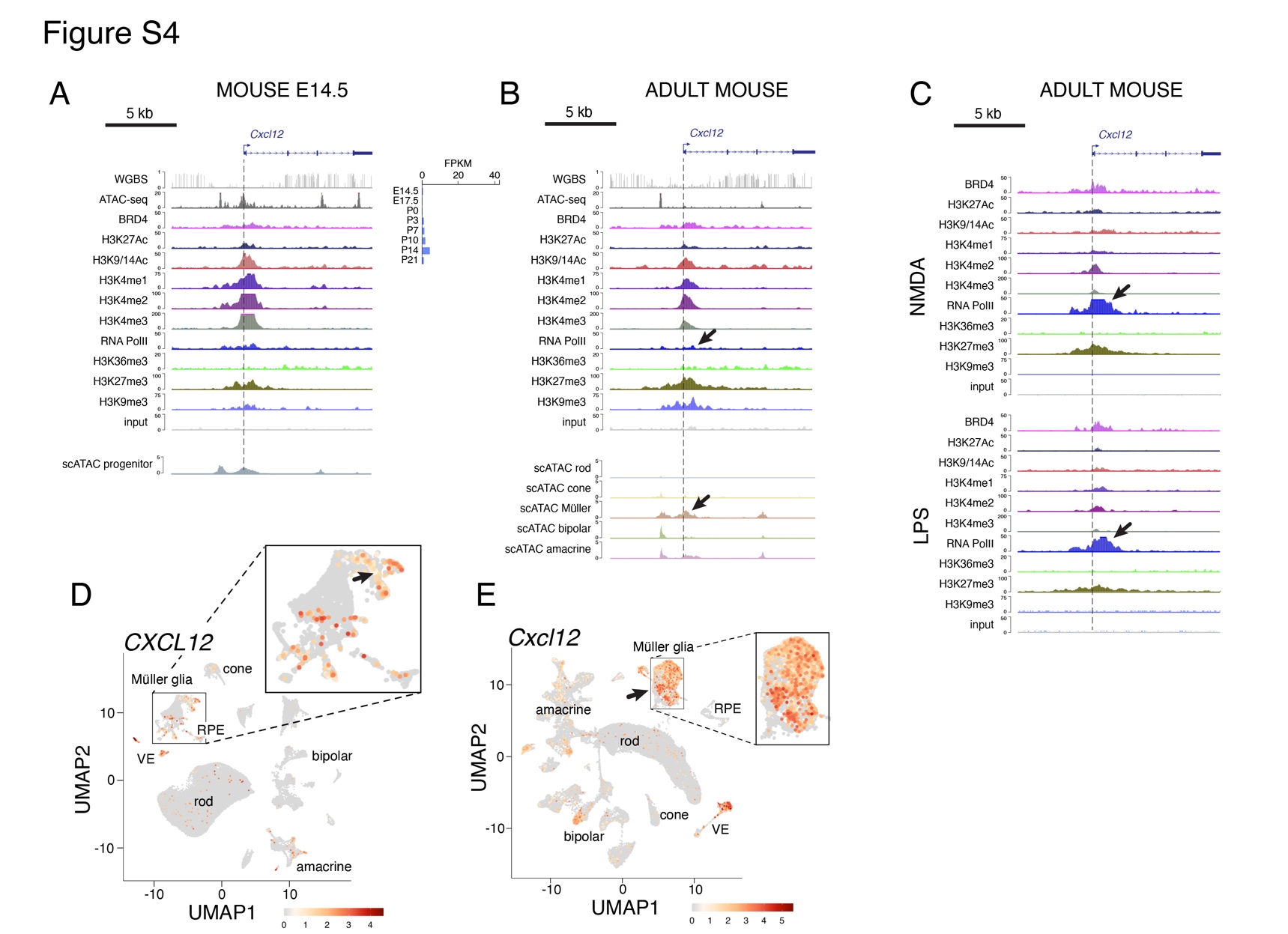

**Figure S5. Induction of specific chemokines and cytokines following stress, injury or disease perturbation. A)** Representative blots showing the subtle increase in cytokine and chemokine levels after 24 hours in culture. This is why we used the untreated cultured retina as a control rather than native retinae. The lower panels show changes following LPS, 40°C and pH7.7 highlighting the differences across conditions. **B)** While the blots are not quantitative, we binned the normalized levels into 4 categories (<1% of background, 1-24% of background, 25-49% of background, >50% of background). Data for all cytokines and chemokines are shown for all stresses. Abbreviations: hr, hours; LPS, lipopolysaccharide; IFN, interferon.

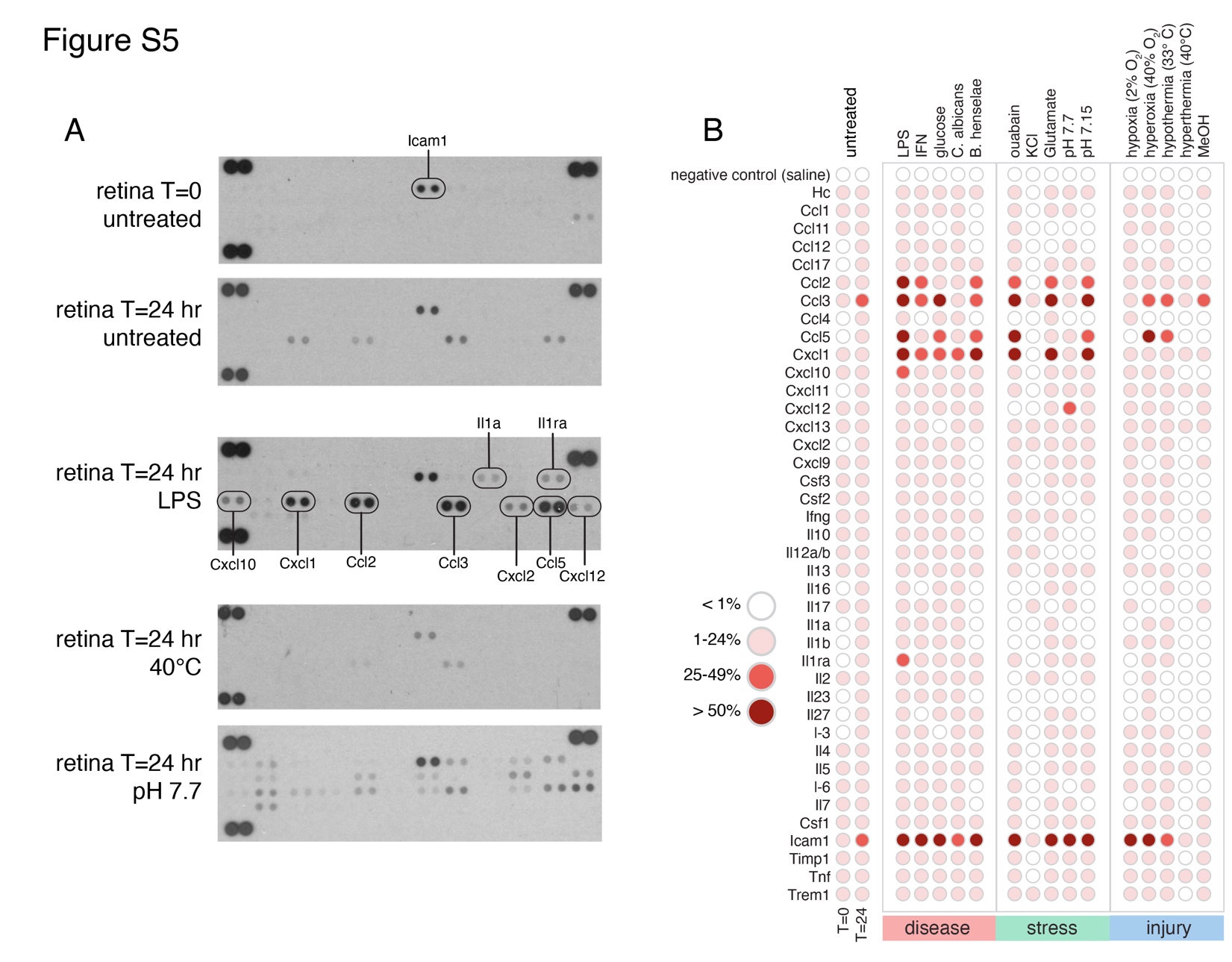

**Figure S6. Zebrafish chemokines have an anti-inflammatory pattern after injury. A)** Barplot showing log2 fold change in expression for the anti-inflammatory (Cxcl18b) and pro-inflammatory (Cxcl12b) genes following NMDA exposure. **B)** Barplot showing log2 fold change in expression for the anti-inflammatory (Cxcl18b) and pro-inflammatory (Cxcl12b) genes following light damage. Data are from Hoang et al. Abbreviations: FDR, false discovery rate.

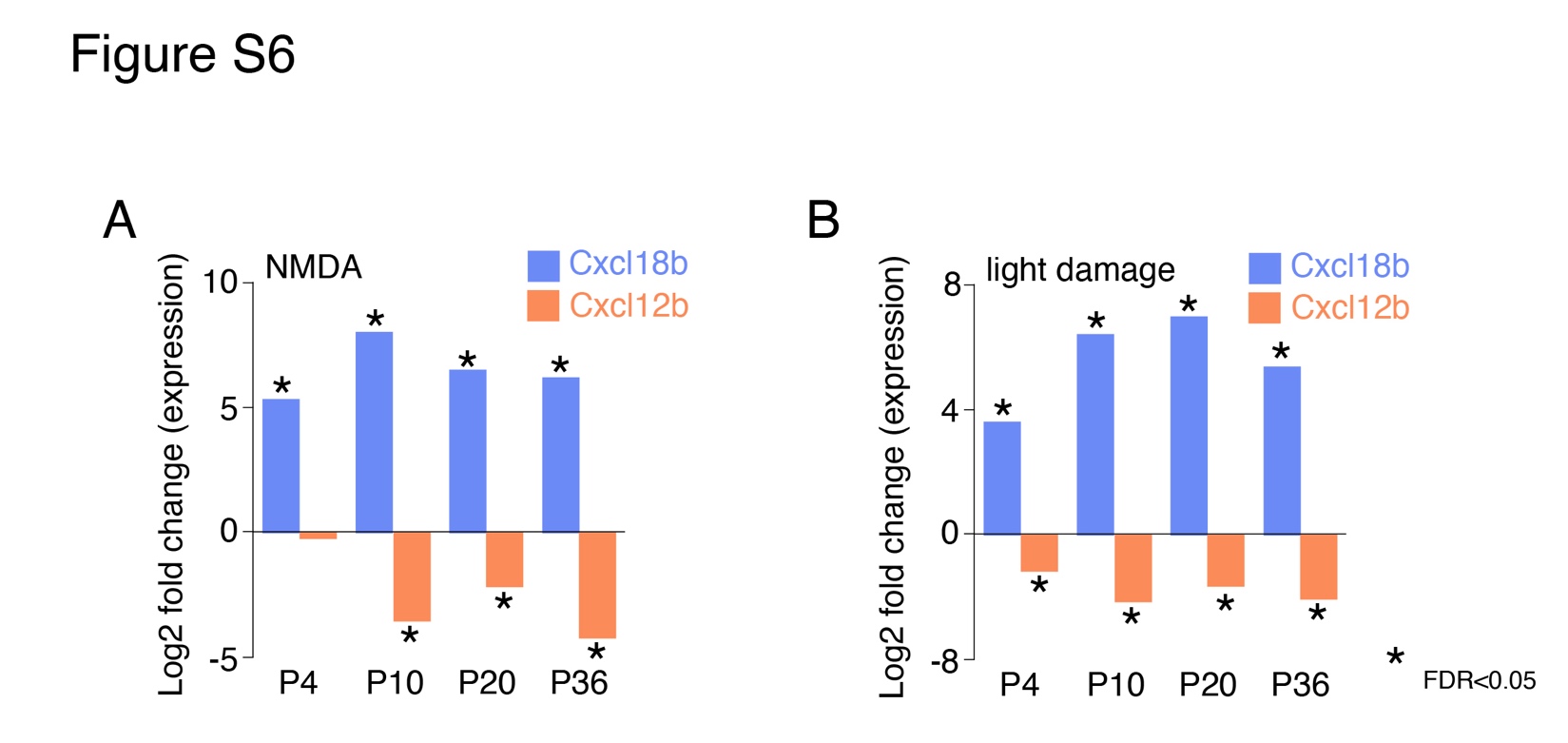

**Figure S7. Pliancy genes are regulated by the transcriptional modules involved in Müller glia reactivity. A)** Tracks for DNA methylation, ATAC-seq and ChIP-seq for *Lhx2* gene in retinal progenitor cells at E14.5. Expression during retinal development is shown in the upper right and scATAC-seq for retinal progenitor cells is shown in the lower panel. *Lhx2* is expressed in retinal progenitor cells and then silenced in the adult retina. **B)** DNA methylation, ATAC-seq and ChIP-seq data are shown for *Lhx2* in the adult retina along with scATAC-seq in the lower panel. The *Lhx2* gene is a Müller glial cell pliancy gene. **C)** DNA methylation, ATAC-seq and ChIP-seq data are shown for the *Lhx2* gene following NMDA and LPS exposure. There is an increase in RNA-PolII binding at the promoter 24 hours after NMDA or LPS exposure consistent with increase in gene expression. **D)** Murine Müller glial cell transcriptional modules from Hoang et al. showing *Lhx2* in the Müller glial cell rest module (M2). **E)** Expression of the *Lhx2* target gene and the transcription factors predicted to activate (Jund (M1), Nr2e1 and Tcf7l2 (M2)) or repress expression (Atf3 (M6), Ets2, Maff (M7)). Abbreviations: kb, kilobase; WGBS, whole genome bisulfite sequencing; FPKM, fragments per kilobase per million reads; LPS, lipopolysaccharide; NMDA, N-methyl-D-aspartate; P indicates promoter proximal binding prediction for the transcription factor.

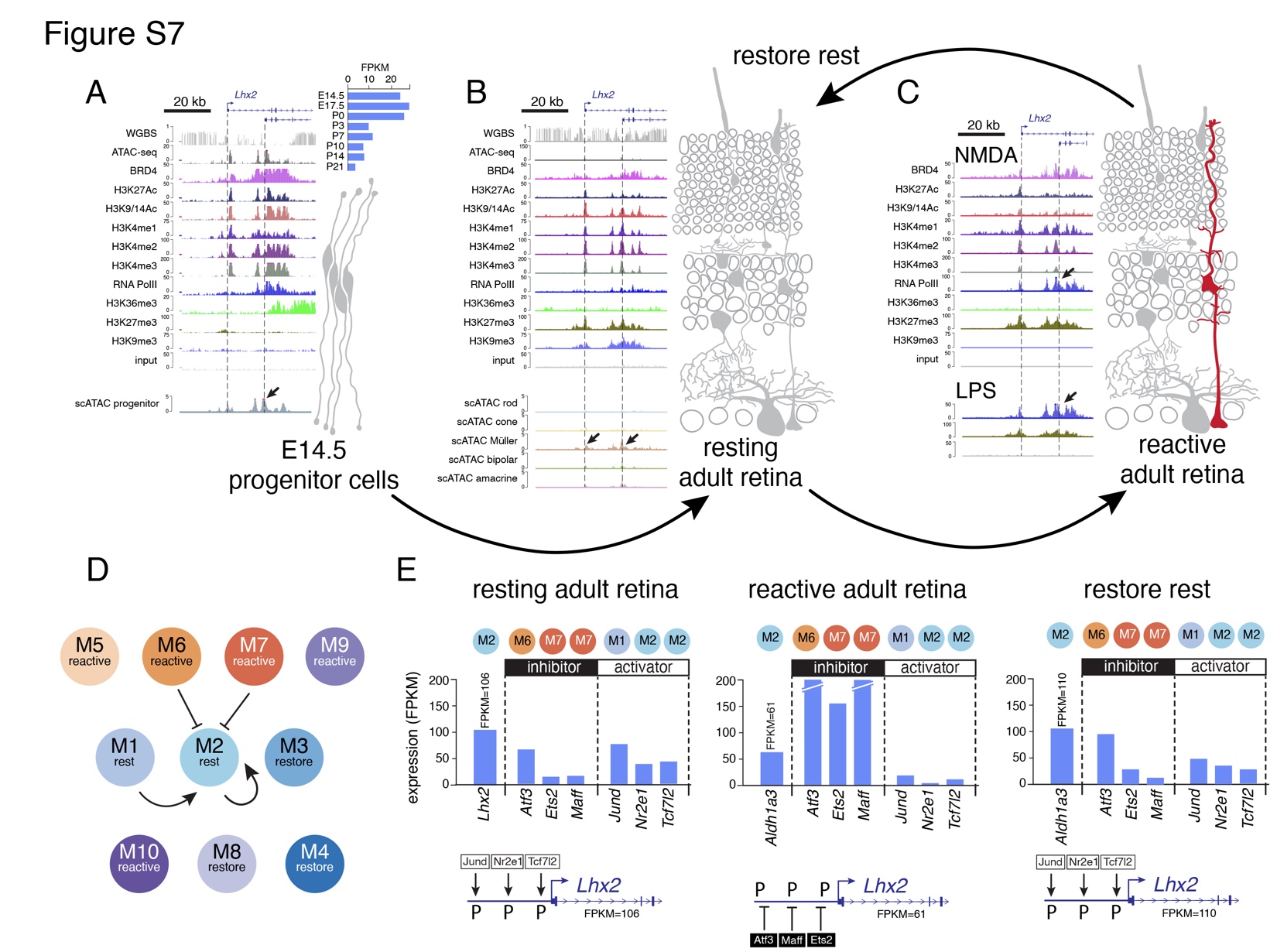

**Figure S8. Pliancy genes are regulated by the transcriptional modules involved in Müller glia reactivity. A)** Tracks for DNA methylation, ATAC-seq and ChIP-seq for *Nfib* gene in retinal progenitor cells at E14.5. Expression during retinal development is shown in the upper right and scATAC-seq for retinal progenitor cells is shown in the lower panel. *Nfib* is expressed in retinal progenitor cells and then silenced in the adult retina. **B)** DNA methylation, ATAC-seq and ChIP-seq data are shown for *Nfib* in the adult retina along with scATAC-seq in the lower panel. The *Nfib* gene is a Müller glial cell pliancy gene. **C)** DNA methylation, ATAC-seq and ChIP-seq data are shown for the *Nfib* gene following NMDA and LPS exposure. There is an increase in RNA-PolII binding at the promoter 24 hours after NMDA or LPS exposure consistent with increase in gene expression. **D)** Murine Müller glial cell transcriptional modules from Hoang et al. showing *Nfib* in the Müller glial cell restore module (M3). **E)** Expression of the *Lhx2* target gene and the transcription factors predicted to activate (Dpp (M2), Plag1 (M8)) or repress expression (Rel, Rela, Nfkb1 (M7), Relb, Nfkb2 (M6)). Abbreviations: kb, kilobase; WGBS, whole genome bisulfite sequencing; FPKM, fragments per kilobase per million reads; LPS, lipopolysaccharide; NMDA, N-methyl-D-aspartate; P indicates promoter proximal binding prediction for the transcription factor.

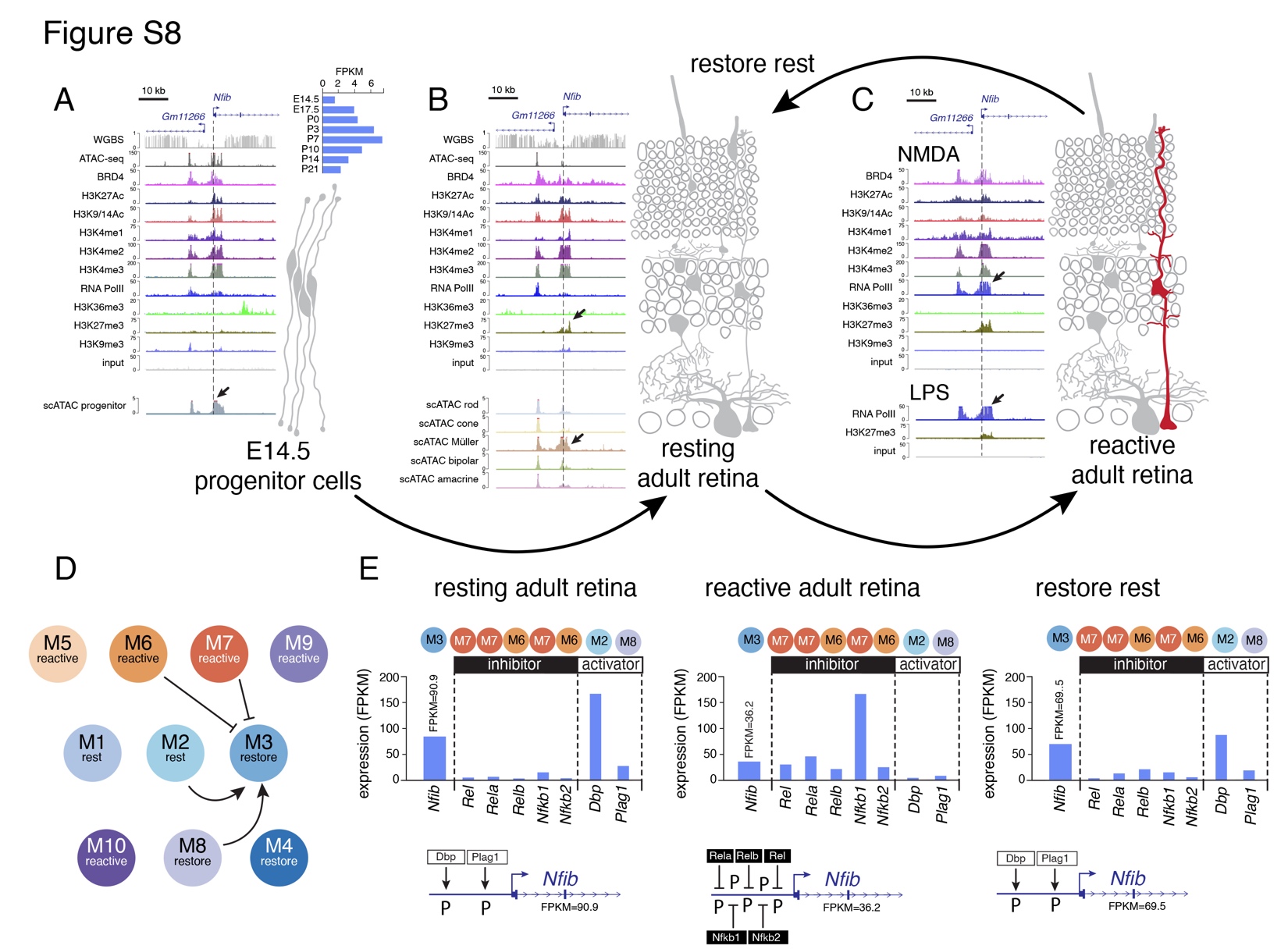

**Figure S9. There are two patterns of chromatin accessibility for the murine orthologues of the zebrafish genes involved in Müller glial cell regeneration. A-C)** DNA methylation (WGBS), bulk ATAC-seq, ChIP-seq and scATAC-seq for E14.5 retina, normal adult retina and adult retina exposed to NMDA for a representative gene (*Kif11)* of the constitutive class of genes. This class includes the differentially expressed genes in zebrafish following NMDA or light damage that have murine orthologues but were not found to be differentially expressed in mouse retina in Hoang et al. The *Kif11* gene is expressed in retinal progenitor cells but is silenced during differentiation. After exposure to NMDA there is increase in expression **D-F)** Similar data are shown as in A-C but for the 2^nd^ chromatin pattern that is restricted to Müller glia. In this example (*Igf2bp2*), the gene is expressed in retinal progenitor cells and then silenced during retinal differentiation. In the adult retina, the promoter is accessible in Müller glia but polycomb repressed (H3K27me3) in other cell types. Following NMDA exposure, the polycomb repression is maintained but RNA-polII binding is increased in correlation with increased expression. Abbreviations: kb, kilobase; WGBS, whole genome bisulfite sequencing; FPKM, fragments per kilobase per million reads; NMDA, N-methyl-D-aspartate.

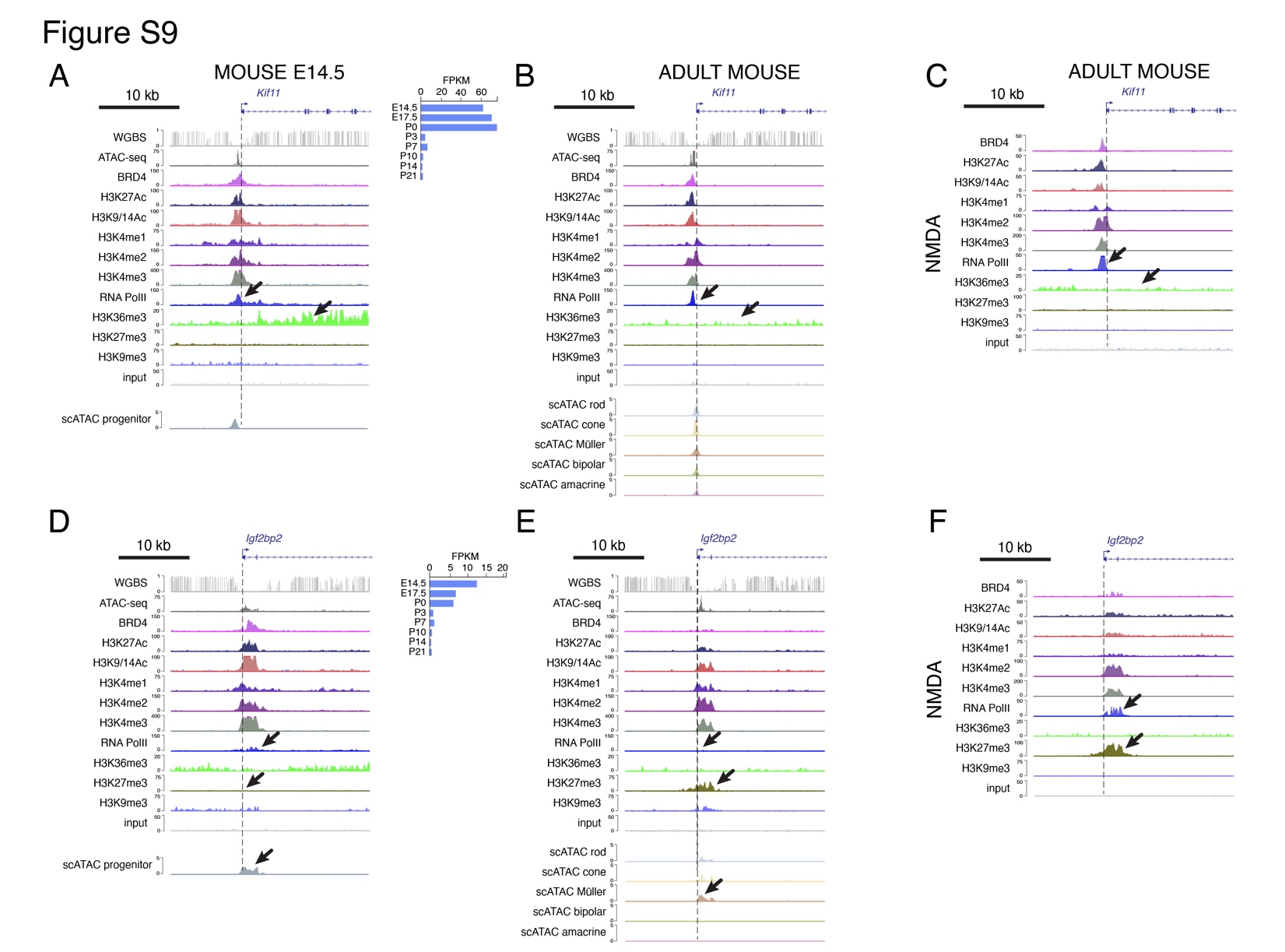

**Figure S10. Sox9, Myb and Ctsc represent a cellular pliancy gene regulatory module. A-D)** DNA methylation (WGBS), bulk ATAC-seq, ChIP-seq and scATAC-seq for E14.5 retina, normal adult retina and adult retina exposed to NMDA for a representative gene (*Ctsc)* of the restricted class of genes. This class includes the differentially expressed genes in zebrafish following NMDA or light damage that have murine orthologues but were not found to be differentially expressed in mouse retina in Hoang et al. The *Ctsc* gene is a pliancy gene (arrows) with accessible chromatin in Müller glia and H3K27me3 repression in other cell types. It is upregulated specifically in Müller glia (D) following stress, injury or disease. **E-H)** DNA methylation (WGBS), bulk ATAC-seq, ChIP-seq and scATAC-seq for E14.5 retina, normal adult retina and adult retina exposed to NMDA for a representative gene (*Sox9)* of the restricted class of genes that is predicted to regulate *Ctsc*. The *Sox9* gene is a pliancy gene (arrows) with accessible chromatin in Müller glia and H3K27me3 repression in other cell types. It is upregulated specifically in Müller glia following stress, injury or disease (H). Abbreviations: kb, kilobase; WGBS, whole genome bisulfite sequencing; FPKM, fragments per kilobase per million reads; NMDA, N-methyl-D-aspartate.

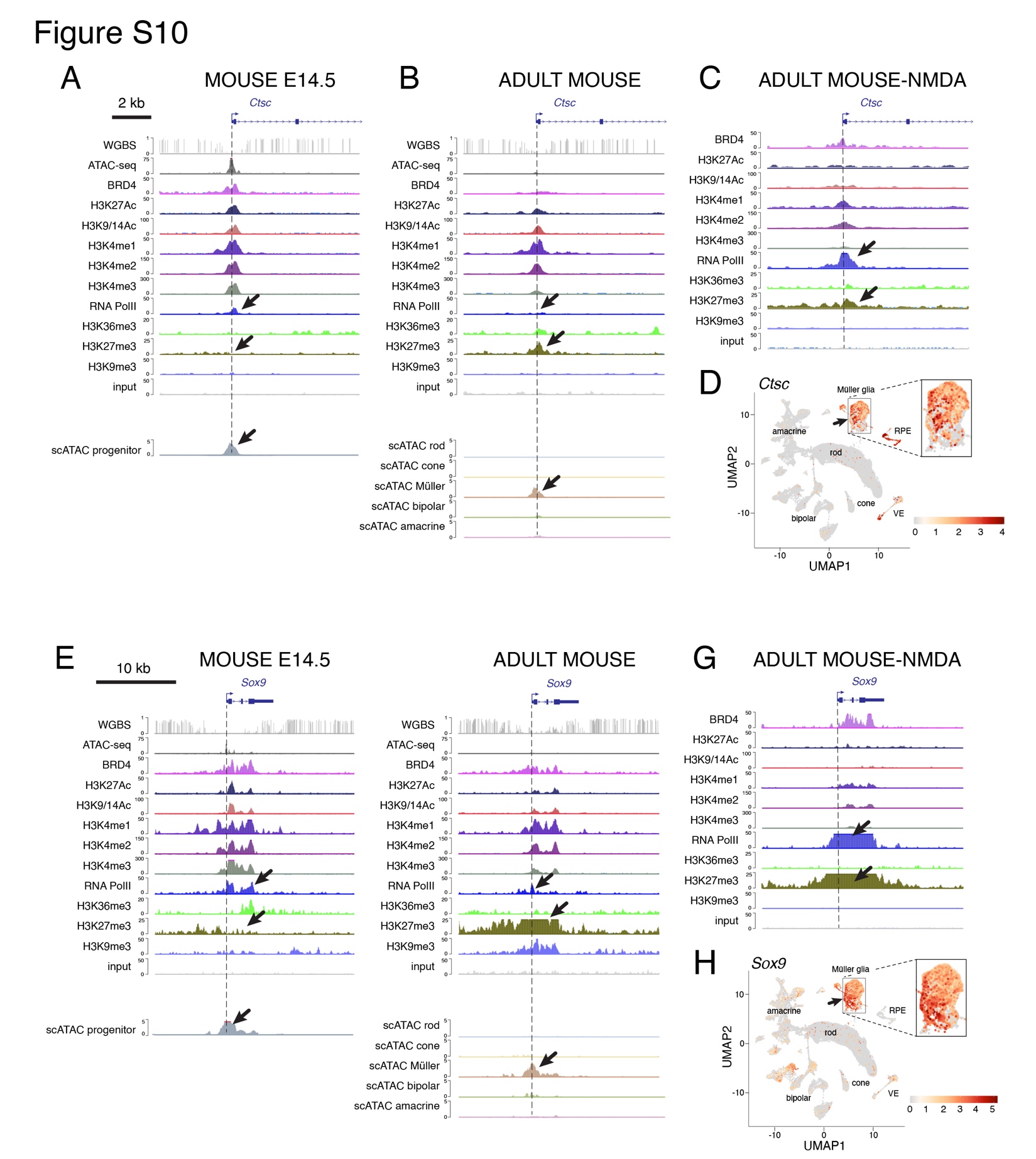

**Figure S11. A rare subset of genes for the murine orthologues of the zebrafish genes involved in Müller glial cell regeneration are constitutively repressed. A,B)** Expression of Prss16 following light damage or NMDA in murine and zebrafish retinae from Hoang et al. **C-E)** DNA methylation (WGBS), bulk ATAC-seq, ChIP-seq and scATAC-seq for E14.5 retina, normal adult retina and adult retina exposed to NMDA for *Prss16* showing constitutive heterochromatin under all conditions. Abbreviations: kb, kilobase; WGBS, whole genome bisulfite sequencing; FPKM, fragments per kilobase per million reads; NMDA, N-methyl-D-aspartate.

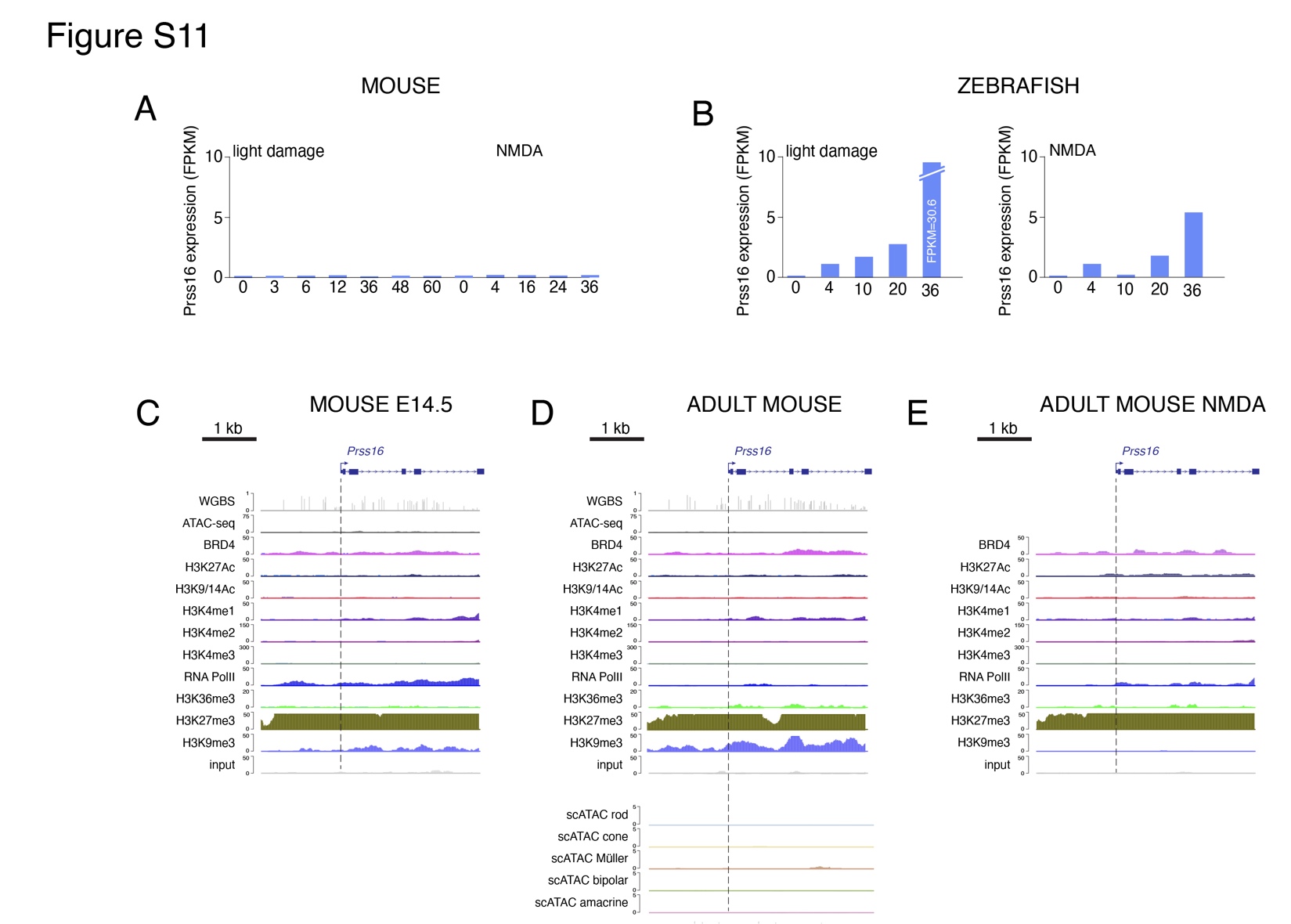
